# Benchmarking the identification of a single degraded protein to explore optimal search strategies for ancient proteins

**DOI:** 10.1101/2023.12.15.571577

**Authors:** Ismael Rodriguez Palomo, Bharath Nair, Yun Chiang, Joannes Dekker, Benjamin Dartigues, Meaghan Mackie, Miranda Evans, Ruairidh Macleod, Jesper V. Olsen, Matthew J. Collins

## Abstract

Palaeoproteomics is a rapidly evolving discipline, and practitioners are constantly developing novel strategies for the analyses and interpretations of complex, degraded protein mixtures. The community has also established standards of good practice to interrogate our data. However, there is a lack of a systematic exploration of how these affect the identification of peptides, post-translational modifications (PTMs), proteins and their significance (through the False Discovery Rate) and correctness. We systematically investigated the performance of a wide range of sequencing tools and search engines in a controlled system: the experimental degradation of the single purified bovine β-lactoglobulin (BLG), heated at 95 °C and pH 7 for 0, 4 and 128 days. We target BLG since it is one of the most robust and ubiquitous proteins in the archaeological record. We tested different reference database choices, a targeted dairy protein one, and the whole bovine proteome and the three digestion options (tryptic-, semi-tryptic- and non-specific searches), in order to evaluate the effects of search space and the identification of peptides. We also explored alternative strategies, including open search that allows for the global identification of PTMs based upon wide precursor mass tolerance and *de novo* sequencing to boost sequence coverage. We analysed the samples using Mascot, MaxQuant, Metamorpheus, pFind, Fragpipe and DeNovoGUI (pepNovo+, DirecTag, Novor), benchmarked these tools and discuss the optimal strategy for the characterisation of ancient proteins. We also studied physicochemical properties of the BLG that correlate with bias in the identification coverage.

## Introduction

Proteomics software has been developed to identify and quantify proteins for different purposes and in various contexts, by processing and analysing mass spectrometry data (Aebersold & Mann, 2003). Many of these tools have been developed to analyse proteins in complex mixtures and map for post-translational modifications (PTMs). Most studies involve querying the mass spectra against a database of possible proteins (database search*, DBs*). Others aim at reconstructing the peptide sequences *de novo* from the spectra, without using any external data (*de novo* sequencing, *dnS*). Increasingly these methods are being applied beyond ‘typical’ tissues for which these software are optimised.

*DBs* approaches work effectively on modern tissues from well-studied organisms with complete sequence databases, typically matching up to 50% of MS/MS queries to peptides (peptide-spectrum matches, PSMs) in a typical tandem mass spectrometry experiment (Cox & Mann, 2008). Larger studies looking at PRIDE (Perez-Riverol et al., 2021) and MassIVE databases have determined that around 75% of submitted spectra remain unidentified (Griss et al., 2016; Bittremieux et al., 2022). Metaproteomics studies attempt to map proteins derived from complex samples such as microbiomes, in which it may be difficult to identify all the targets present without a proteogenomics strategy. The same is true for highly variable sequences, such as those found in immunoglobulins or novel sequences as found in cancers (Alfaro et al., 2017). *dnS* tools like PEAKS (Zhang et al., 2012) were initially developed to characterise these sequences. In addition to novel proteins introduced from the microbial community, the application of proteomics to food and forensics is complicated by an increase in the number of PTMs due to both enzymatic and chemical modifications or introduced by bacterial effectors (Liu et al., 2023).

Palaeoproteomics, the study of ancient proteins that are often fragmented and heavily modified, combines all of the above challenges. From studies of historical artworks to wider studies of material culture or skeletal remains in archaeology and palaeontology, chemical degradation adds further to the challenges of identification. In the field, most published ancient spectra remain unidentified. *De novo* sequencing has been decisive in studies looking at the phylogeny and evolutionary history of extinct organisms (Welker et al., 2015, 2019; Chen et al., 2019; Presslee et al., 2019; Cappellini et al., 2019). However, noise and missing ions might produce errors that can never be identified without matching to a database sequence. Muth and Renard (2018) evaluated the accuracy of different *dnS* software. They showed that only around 35% of the complete peptide sequences were correct on real HCD (higher-energy collisional dissociation) data and 85% on simulated data. On a different front, there are increasing efforts to minimise and identify modern protein contaminants in ancient samples. Deamidation rates have been proposed and applied to authenticate ancient proteins (Ramsøe et al., 2020, 2021). However, a clear correlation between deamidation rates and time is yet to be established (Schroeter & Cleland, 2016). A deeper understanding of deamidation and other degradation patterns is key for ancient protein authentication.

At present, the field of palaeoproteomics has tended to use the same software tools as those originally designed for identification on targeted tissues. And due to the awareness of the specific challenges, the community sets and demands stringent approaches to interrogate the data (Hendy, Welker, et al., 2018; Hendy, Warinner, et al., 2018; Warinner et al., 2022). This includes the consideration of an increasingly wide array of post-translational modifications (PTMs), well-represented databases which encompass every potential constituent of samples, robust rules on the identification of peptides and proteins and statistically significant false discovery rates (FDRs) (Benjamini & Hochberg, 1995; Choi & Nesvizhskii, 2008; Käll et al., 2008). PSMs leading to extraordinary claims or novel peptides are manually checked, and at least 2 uniquely identified peptides are required to claim the presence of a specific protein. However, there is a lack of a systematic assessment of the effectiveness of these tools, approaches and standards in identifying ancient degraded proteins. The number of new strategies, software and packages being continuously developed to address problems with *dnS* and *DBs* already highlights issues with the different strategies. Because of all this, there is an increasing demand for all raw data to be made public in proteomics repositories for reanalysis. Along with it, the use of free and open-source software, standards and formats facilitates reproducibility and the adoption of common practices.

When using *DBs* software to analyse mass spectrometry data obtained from ancient protein samples, researchers often face decisions concerning parameter selection tailored to the specific sample types. These choices require balancing compromises and relying on a clear research question and hypothesis, and expert, multidisciplinary knowledge about the context of the samples. Broadly, one needs to consider the size of the search space, significance of results, computational resources and time. If the search space is too large, the memory usage and computation time increase and the significance of the identifications is compromised (Jeong et al., 2012; Noble, 2015). On the other hand, if the search space is too small, there is the risk of missing proteins present in the samples. In *de novo* sequencing, the peptide sequence is derived directly from the spectra, and therefore, it frees researchers from these decisions and compromises. However, care must be taken with the accuracy of the sequences (Muth & Renard, 2018; Beslic et al., 2023), especially in phylogenetic and evolutionary studies dealing with novel peptides of extinct, unsequenced species. In both *DBs* and *dnS*, peptides of interest are usually validated using BLASTp or tBLASTn (Altschul et al., 1990) against a reference database.

The search parameters that most impact the search space size and need more tuning are the type of enzymatic cleavage, the protein database, and the PTMs targeted. Evaluating the degree of sample degradation prompts the researcher to determine the suitable enzymatic cleavage method (specific, semi-specific or non-specific) and the post-translational modifications (PTMs) to be targeted during the search. The specific archaeological and palaeontological questions that the researcher is trying to answer, along with the type of sample, determine the scope of the database. The size of the search space in turn affects the significance of the results. To control for the significance of multiple PSMs, the false discovery proportion (FDP) of a search is estimated using a Target-Decoy Competition (TDC) (Moore et al., 2002; Elias & Gygi, 2007), from which a False Discovery Rate (FDR) is obtained. A decoy database is built by reversing the sequences in the target database and it competes together for PSMs. It therefore contains known incorrect sequences that share statistical properties with the target database like amino acid distribution or peptide length. In the end, it will result in known incorrect PSMs that would have otherwise been considered positive. This allows sorting PSMs by a matching score and setting filters so that an expected given fraction (the FDR) of PSMs are false (the decoy ones). Thus the FDR is just an expectation of the actual proportion of false discoveries (the FDP). In fact, several studies have pointed out the divergence between the FDP and FDR (He et al., 2015; Madej & Lam, 2023; Ebadi et al., 2023), and the “instability” of the TDC approach that can lead to an underestimation of the FDP in high-resolution mass spectrometry (Couté et al., 2020). Moreover, FDRs are challenging in the study of complex or uncharacterised proteomes, as “treating each search result equally is methodically incorrect when peptides and proteins are not equally likely to be measured by LC-MS/MS and identified by search engines” (Wang et al., 2022).

In this study, we wish to explore which solution is optimal for degraded proteins given these constraints. For this, we create a deliberately simple experimental dataset which focuses not on the diversity of a complex mixture, but on the changes caused by mild and extreme degradation. Our model is a suspension of a small and well-characterised protein, two isoforms of a 156 amino acid long lectin, bovine β-Lactoglobulin (BLG), heated at neutral pH conditions for 0, 4 and 128 days, respectively. The three time points are designed to test the ability of these datasets to identify a mild (trypsin cleaved) and extreme example of protein degradation in which most surviving peptides are amenable to mass spectrometry and enzymatic digestion is not necessary. We explore a range of different software solutions and strategies to identify MS2 queries generated by this experiment and discuss the results in connection with the common practices and challenges in palaeoproteomics.

We chose BLG, as it is one of the most commonly preserved and identified proteins in dental calculus and ceramics and thus has become the main protein used in archaeology to explore the spread of dairying in the past (Warinner et al., 2014; Hendy, Warinner, et al., 2018; Hendy, Colonese, et al., 2018; Jeong et al., 2018; Charlton et al., 2019; Wilkin et al., 2020, 2021; Bleasdale et al., 2021; Tanasi et al., 2021). It is suggested that mineral surfaces play an important role in the survival of this protein (Fonseca et al., 2022), as it has been also found for other eggshell proteins (Demarchi et al., 2016). Evans et al. (2024) explored a range of protein properties that might enable their survival in fresh ingredients and in cooked foodcrusts before and after burial. We explored as well several physico-chemical properties of the BLG: structural flexibility, solvent exposure, amyloidogenicity and isoelectric point; and linked them to the degradation, detection and identification of different regions of the BLG.

## Materials and Methods

### The target protein

BLG was purchased from Sigma Sigma Aldrich (CAS No.: 9045-23-2). It is a single purified protein with purity greater than 90% and traces of other bovine milk proteins. No further purification was undertaken.

### Experimental setup

1.75 mg of BLG was dissolved in 7 mL of Molecular Grade Millipore water at pH ∼7 (to a final concentration of 250 μg/mL) in borosilicate glass vials closed with screw caps with polytetrafluoroethylene (PTFE) lining on the inside.

Samples and blanks were placed on a VWR^®^ dry block heater with a heated lid maintained at a constant temperature of 95 °C for 128 days. 200 μL was sampled from each glass vial at designated time lengths (0, 4 d, and 128 days) to 1 mL Eppendorf Safe-Lock^®^ tubes. Samples were then subjected to vacuum centrifuge at 60 °C to dry the thermal degradation products and were stored at -20 °C. A USB data logger (ebro EBI 310-T1) connected to an external sensor was used for monitoring temperature throughout the experiment. A schematic representation of the experimental setup is shown in Figure 1a.

**Figure 1.**
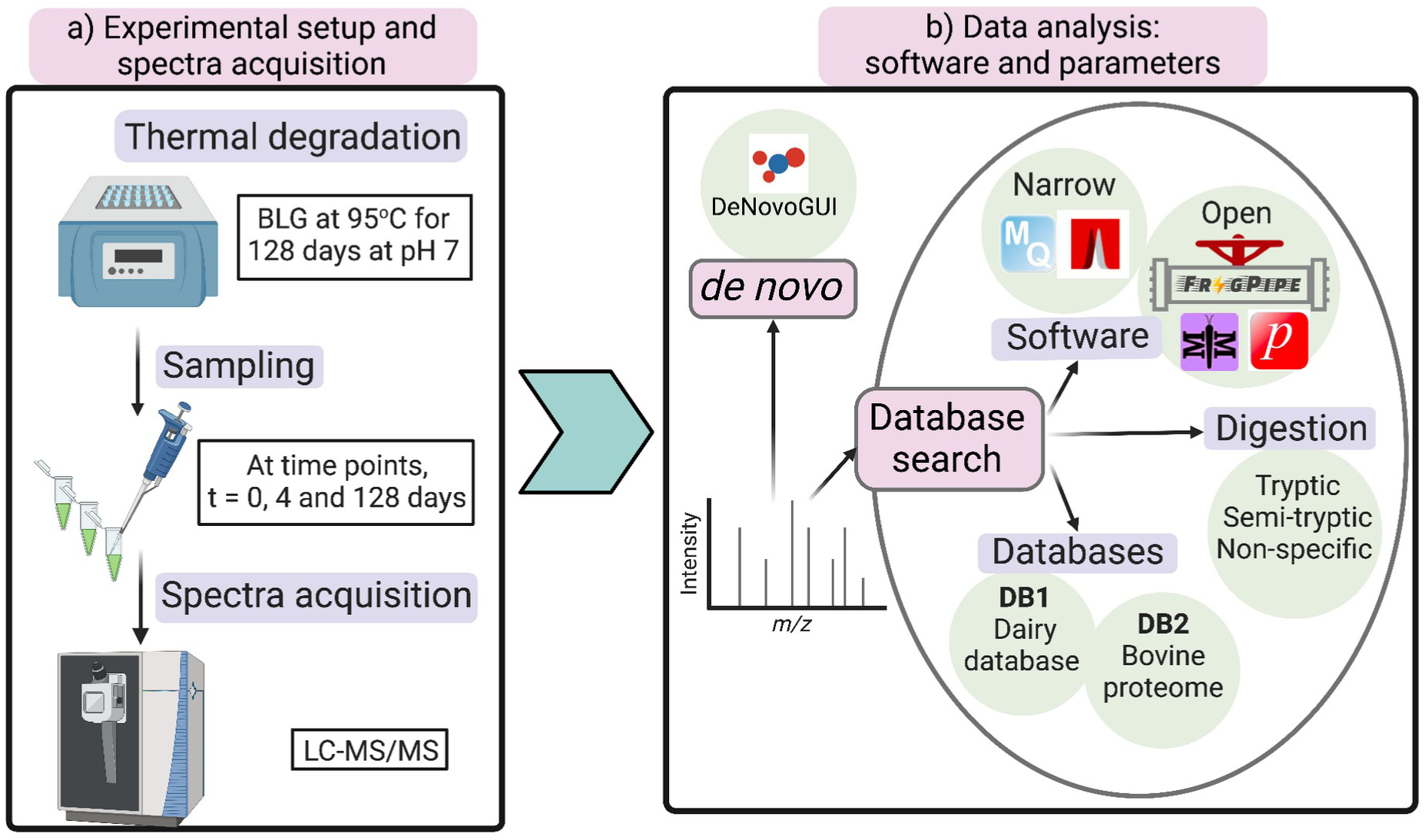
Schematic representation of the experimental setup (a) and the peptide identification strategies tested, including the different software, databases, and digestion (b).

### LC-MS/MS

The peptides from the 0 and 4-day experiments were extracted following a previously published protocol (Cappellini et al., 2019) for degraded samples that consist of denaturation, reduction, alkylation, enzyme digestion, and desalting as described below.. Samples from each glass vial were resuspended to 300 μL solution consisting of 2 M guanidine hydrochloride (GuHCl, Thermo Scientific), 100 mMTris buffer (pH ∼8), 20 mM 2-chloroacetamide (CAA) and 10 mM Tris (2-carboxyethyl)phosphine (TCEP) in ultrapure water. 0.2 μg of mass-spectrometry grade rLysC (Promega) enzyme was added prior to incubation with shaking for 2 h at 37 °C. Samples were subsequently diluted to 0.6 M GuHCl and 0.4 μg of mass-spectrometry grade trypsin was added and subjected to overnight incubation with shaking at 37 °C. The peptides at 128 days were alkylated but not digested by trypsin.

On the next day, samples were acidified using 10 % *(v/v)* trifluoroacetic acid (TFA) and purified using in-house prepared C18 solid-phase extraction StageTips™ (Rappsilber et al., 2007) by following a previously published protocol (Cappellini et al., 2019) described as follows. Stage-Tips were prepared in-house wherein two disks, pre-pierced with a 14 gauge blunt-end needle, were pushed out of an Empore C18 membrane (3M) into the end of each P200 micropipette tip. The Stage-Tips were sequentially conditioned with 150 μL of methanol, 150 μL of 80% acetonitrile solution (containing 80% acetonitrile, 0.1% trifluoroacetic acid (TFA), and ultrapure molecular biology-grade H2O -v/v/v-), and 150 μL of 0.1% TFA in H2O (v/v). Acidified peptides were then loaded onto the Stage-Tips and were immobilized on the C18 filter by centrifugation at 1300 ×g until all the solvent had passed through. The filter was washed with 150 μL of 0.1% TFA in H2O (v/v), was centrifuged at 1300 ×g until dry. Immobilized peptides within Stage-Tips were eluted using 150 μL of 50 % acetonitrile solution (consisting of 50 % acetonitrile, 0.1 % TFA, and ultrapure molecular grade water) and was stored at -20°C.20 to 25 μL of eluted samples were transferred to the MS analysis facility at the Novo Nordisk Foundation Center for Protein Research (CPR), University of Copenhagen. Samples were subjected to nanoflow liquid chromatography-tandem mass spectrometry (nano-LC MS/MS) using an in-house packed column on an EASY-nLC ™ 1200 system connected to an Orbitrap Exploris 480 (Thermo Scientific, Germany) mass spectrometer. A column temperature of 40 °C was maintained with an integrated column oven. For each run, 3 uL of sample was separated using a linear gradient from 5% to 30% buffer B in 25 minutes followed by a 2-minute step to 45% buffer B. Following the linear gradient, the column was washed by increasing the concentration of buffer B to 80% in 2 minutes, and remaining there for 2 minutes. Finally, it was re-equilibrated back to 5% in 2 minutes and held for 2 minutes, resulting in a final acquisition time of 35 minutes.

The Orbitrap Exploris 480 was operated in data-dependent acquisition mode using a Top 10 method. The spray voltage was at 2 kV, the S-lens RF level was at 40%, and the ion transfer tube was kept at 275 °C. Full scan MS were acquired at a resolution of 120,000 in a mass range of 350-1400 with an Automatic Gain Control (AGC) target of 300 and a maximum ion injection time of 25 ms. Fragment MS/MS spectra were recorded using Higher-energy Collisional Dissociation (HCD) with a maximum ion injection time set to 118 ms with an AGC target value of 200 at a resolution of 60,000 with a fixed first mass of 100 *m/z*. Normalised collision energy (NCE) was 30 %. The isolation window was set to 1.2 *m/z*, and the dynamic exclusion was 20 seconds.

### Search strategies

#### Open and narrow database search software

We chose various software based on different principles and algorithms (Table 1 and Figure 1b). We chose a range of free, open-source or closed-source software, but included Mascot, which is licensed, since it is frequently used in palaeoproteomics.

**Table 1.** Software used for benchmarking.

| Table 1. Software used for benchmarking |  |  |  |
| --- | --- | --- | --- |
| Software | Type of search | Platform | Citation |
| Fragpipe v19.1 with MSFragger v3.7 | DBs open | University of Cambridge HPC<sup>1</sup>. 56 CPUs in Intel Cascade Lake node and 3.5GB per CPU | Kong et al., 2017; Chang et al., 2020; da Veiga Leprevost et al., 2020; Geiszler et al., 2021 |
|  | DBs narrow |  |  |
| Open-pFind 3 v3.2.0 | DBs open | MiniMax<sup>2</sup>. 24 AMD Ryzen Threadripper 3960X CPUs and 64 GB RAM | Chi, Liu, Yang, Zeng, Wu, Zhou, Niu, et al., 2018 |
| Metamorpheus v0.0.320 | DBs Multi-notch | Mjolnir<sup>3</sup>. 20 CPUs in AMD EPYC 4773 node and 20GB CPU | Solntsev et al., 2018 |
| MaxQuant v2.1.0.0 | DBs narrow | MiniMax<sup>2</sup>. 24 AMD Ryzen Threadripper 3960X CPUs and 64GB RAM. | Tyanova et al., 2016 |
|  |  | University of Cambridge HPC<sup>1</sup>. 56 CPUs in Intel Cascade Lake node and 3.5GB per CPU<sup>+</sup> |  |
| Mascot v2.8.2 | DBs narrow | Mascot Server<sup>4</sup> | Perkins et al., 1999 |
| Novor v1.05.0573 | dnS | MiniMax<sup>2</sup> with DeNovoGUI (Muth et al., 2014). 24 AMD Ryzen Threadripper 3960X CPUs and 64GB RAM | Ma, 2015 |
| DirecTag v1.4.66 | dnS hybrid |  | Tabb et al., 2008 |
| PepNovo+ v3.1 | dnS |  | Frank & Pevzner, 2005; Frank, 2009a; b |

The LC-MS/MS files were run in Thermo RAW file format on pFind3, Metamorpheus, MaxQuant and Fragpipe. For Mascot and DeNovoGUI the files were transformed to MGF (Mascot Generic Format) using msconvert (Chambers et al., 2012).

*DBs* software can be categorised based on the approach for the identification of PTMs and the precursor mass tolerance. In narrow-search, the targeted PTMs are defined *a priori,* and predicted peptide mass including PTMs must match the measured precursor mass with a narrow error. In contrast, open search allows wide precursor mass tolerances in order to identify peptides with unaccounted PTMs “*on-the-fly*”. Multi-notch sets a specific set of precursor mass shifts instead of a range. We note that most narrow-search software (e.g. Mascot, MaxQuant) can in theory also perform open search, by just establishing a large precursor mass tolerance. Unlike specifically authored tools (e.g. pFind, Fragpipe or Metamorpheus), such software lacks other key components for open search. In Fragpipe, MSFragger implements a localization-aware *open search* and the fragment index allows a fast search (Kong et al., 2017; Yu et al., 2020), while Crystal-C (Chang et al., 2020) discards artefacts, and PTMshepherd (Geiszler et al., 2021) helps summarising and analysing open search results. pFind is also optimised for fast searches and is also able to localise PTM mass shifts (Chi, Liu, Yang, Zeng, Wu, Zhou, Niu, et al., 2018). Metamorpheus uses the G-PTM-D workflow, which runs a multi-notch search and an augmented narrow database search to speed up the process (Solntsev et al., 2018). We will refer to Metamorpheus as an open or multi-notch search software.

Conversely, open search software also runs in narrow or closed search mode. Fragpipe was run both in narrow and open mode, referred to as narrow-Fragpipe and open-Fragpipe. For both the open and narrow searches PeptideProphet (Keller et al., 2002; da Veiga Leprevost et al., 2020)) was run for the validation step. Percolator is not compatible with open search. pFind was only run in Open search mode, and it will be referred to as Open-pFind or pFind.

Each software was run using 6 different combinations of parameters (Table 2) that broadly change the search space. This can be appreciated in the number of peptides to search in the spectra that each one generates (Table 2). The selection of parameters is explained in the sections below.

**Table 2.** Database Search Parameters.

| Run # | Database | Enzymatic cleavage | # peptides |
| --- | --- | --- | --- |
| 1 | DB1: dairy database<br>227 proteins and 62,185 residues | Tryptic | 9,302 |
| 2 |  | Semi-tryptic | 102,127 |
| 3 |  | Non-specific | 1,092,998 |
| 4 | DB2: bovine proteome<br>37,606 proteins and 22,906,446 residues | Tryptic | 3,640,393 |
| 5 |  | Semi-tryptic | 38,385,707 |
| 6 |  | Non-specific | 415,906,748 |

#### Dairy database and bovine proteome

Despite the fact that we know that the samples were almost purely composed of BLG and perhaps contaminants, we used two of the widely used database strategies, in order to provide robust and comparable statistics with other studies. Broadly, we define a wide database as one that covers a limited set of proteins from a wide range of species. Many studies use a wide, targeted database, for example (Cappellini et al., 2019; Evans et al., 2023) as they look for just specific proteins (dairy or enamel) from different species. On the opposite side, a deep database is one that represents the whole known proteome or proteomes from a specific species or taxa. These are used, for example, in medical studies that focus on a specific organism, like a human, mouse or zebrafish. But in the study of ancient proteins usually a wide and deep strategy is adopted. A taxonomically broader database is used to explore the wider metaproteome, or dietary proteins with the entire SwissProt and TrEMBL (c. 251 million protein entries) (Bairoch & Apweiler, 2000) being the most typical target (Warinner et al., 2014; Wilkin et al., 2020, 2021; Bleasdale et al., 2021) although even larger ones are used (Jersie-Christensen et al., 2018).

Here we tested the two opposite types of databases: a wide one from (Evans et al., 2023) that includes common and reviewed dairy proteins (DB1) and as a deep one, the Bovine proteome (DB2) (Table 2 and Figure 1b). Both include common contaminants usually reported in palaeoproteomics (Evans et al., 2023) and 4 different bovine BLG variants. DB1 comprises 227 protein entries, while DB2 contains 37,606 proteins.

All software were run with both databases, except for narrow-Fragpipe, which was run only on the DB1.

#### Digestion: tryptic, semi-tryptic and non-specific

Most proteomics protocols include a digestion step, usually with trypsin. However, in palaeoproteomics the most degraded samples require no digestion step, removing the challenge of untangling diagenesis from the proteolytic background (Demarchi et al., 2016). This is the case with the 128-days sample. To test the differences in performance between using a tryptic, semi-tryptic and non-specific search we applied all three approaches on the three samples, each with an increasing degree of degradation (Table 2 and Figure 1). This is then reflected in the database search, which generates tryptic peptides (the two ends are tryptic, ie. result from a tryptic cleavage) out of the selected database to match the sample. However, in palaeoproteomics, samples are usually degraded, and hydrolysis generates a non-trivial amount of non-tryptic peptides. This means that the database search will also need to run on semi-tryptic mode, which generates peptides with just one tryptic end. The non-specific search produces all possible peptides within a certain length range. Semi-specific and especially non-specific searches increase the search space dramatically and therefore the running time and memory requirements. If combined with a large database, it might not be feasible to run for some software and systems.

#### Other parameters

We set the rest of the parameters as default or to values commonly used in Palaeoproteomics, to emulate what a standard run would be in a real scenario. We include glutamine and asparagine deamidation, oxidation of methionine, and N-terminal acetylation as variable PTMs, and cysteine carbamidomethylation as fixed. Peptide length range is 7 to 25 amino acids and the peptide mass range is 500 to 5000 Da. We allow up to 2 missed cleavages and turn off the match between runs feature in the software that implements it, so they can be compared and to keep identifications independent between the 3 samples. Supplementary File S1 contains more detailed lists of parameters, compiled or adapted from the parameter files used in the set up of the searches.

### Collation of Database Search (DBs) results

The different software reports results using different formats. These include pepXML, mzIdentML or a tab or comma-delimited table. We used Python 3.9 with lxml, pyteomics (Levitsky et al., 2019) and pandas packages to collate the results into a single PSM data frame. Each row is a single PSM from each sample and run. We processed all the results and extracted the following information from each PSM: Scan #, retention time, peptide sequence, protein accession, start and end position, whether it is a decoy match, calculated mass, delta mass, and q-value. q-value is the minimum FDR threshold at which the identification can be considered statistically significant. We calculated PSM-level q-values using pyteomics *q_value* function. We only include the best PSM per scan. When a peptide is matched against BLG from different species, if the bovine BLG is one of them, the start and end positions of the peptide are reported with respect to that one.

In instances where the same spectra is identified by different software and assigned to BLG, we calculated the Levenshtein distance between the two peptide identifications. This is the number of insertions, deletions and substitutions that transform one sequence into the other. We used the Python package Levenshtein (Bachmann, 2024).

After removing decoy matches and non BLG proteins from the PSM data frame, we pivoted it into a long format, taking each peptide’s amino acid position into a single row. For example, a PSM row with peptide sequence TPEVDDEALEK, which starts at position 141 in the protein, is turned into 11 rows, one per amino acid, and with positions 141, 142, 143, … and 152. When all PSMs are considered, this allows for calculating each position’s coverage along the protein sequence.

In this and subsequent analysis, the BLG amino acid positions will be given with respect to the beginning of the immature sequence, ie. counting the signal peptide (as they are given also in figures 7, 8 and 9). We do this to explicitly acknowledge the existence of the signal peptide, which at the same time is not present in the mature molecule in milk whey, and therefore there should not be any peptide starting before position 17. Moreover, signal peptides are always included in the protein database sequences used in database search engines.

### *De novo* sequencing

These methods use machine learning techniques and employ graphs as data structures and algorithms applied to them. We run Novor, DirecTag and PepNovo+ using DeNovoGui (Muth et al., 2014) interface (Table 1 and Figure 1b). Novor uses two decision trees trained from peptide spectral libraries for its scoring PSM function. DirecTag, follows a hybrid approach and uses a database to extend sequence tags derived from the spectra (Tabb et al., 2008; Zhang et al., 2012). PepNovo+ uses probabilistic network modelling and fragment ion intensity ranks prediction and scoring (Frank & Pevzner, 2005; Frank, 2009a; b) and is followed by a MS-BLAST (Shevchenko et al., 2001) query of the peptides against a database. We ran DeNovoGui with both sequence databases, DB1 and DB2, to understand how they can also affect these hybrid methods. pNovo+, which is included in DeNovoGUI did not produce enough meaningful results and was discarded from further analysis.

### Mapping of *de novo* peptides

We mapped the *de novo* peptides back to the bovine BLG protein sequence allowing mismatches, that in this case account for sequencing errors. Since leucine (L) and isoleucine (I), aspartic (D) and glutamic acid (E) and deamidated asparagine (N) and glutamine (Q) have the same mass respectively, we ‘blind’ all the sequences, so that mismatches between these pairs are not counted as errors. I and L are encoded as a B, D and N as a X and E and Q as Z. Having all sequences blinded, we first create a 3, 4 and 5-mer index from the bovine BLG variants (BLG k-mer index). Then for each peptide, we generate a list of C and N-terminal 3, 4 and 5-mers and matched against the BLG k-mer index. These were used as seed matches that we extended by running a pairwise alignment using the Needleman-Wunsch (Needleman & Wunsch, 1970) algorithm. When a peptide matches to k-mers from different positions, the alignment with the highest identity is kept. In case of a tie, we keep the one originating from a longer k-mer, giving preference to C-terminal k-mers. We allow gaps in the alignment, accounting for errors in the *de novo* sequencing in which an amino acid is skipped or extra ones are added, due to noise in the spectra or missing fragment ions. Finally, alignments with overall identity below 70% are discarded. Figure 2 shows a diagram of the procedure. Once the peptides are mapped we pivot the data into long format as before, so we can calculate the coverage and accuracy of each position along the BLG sequence.

**Figure 2.**
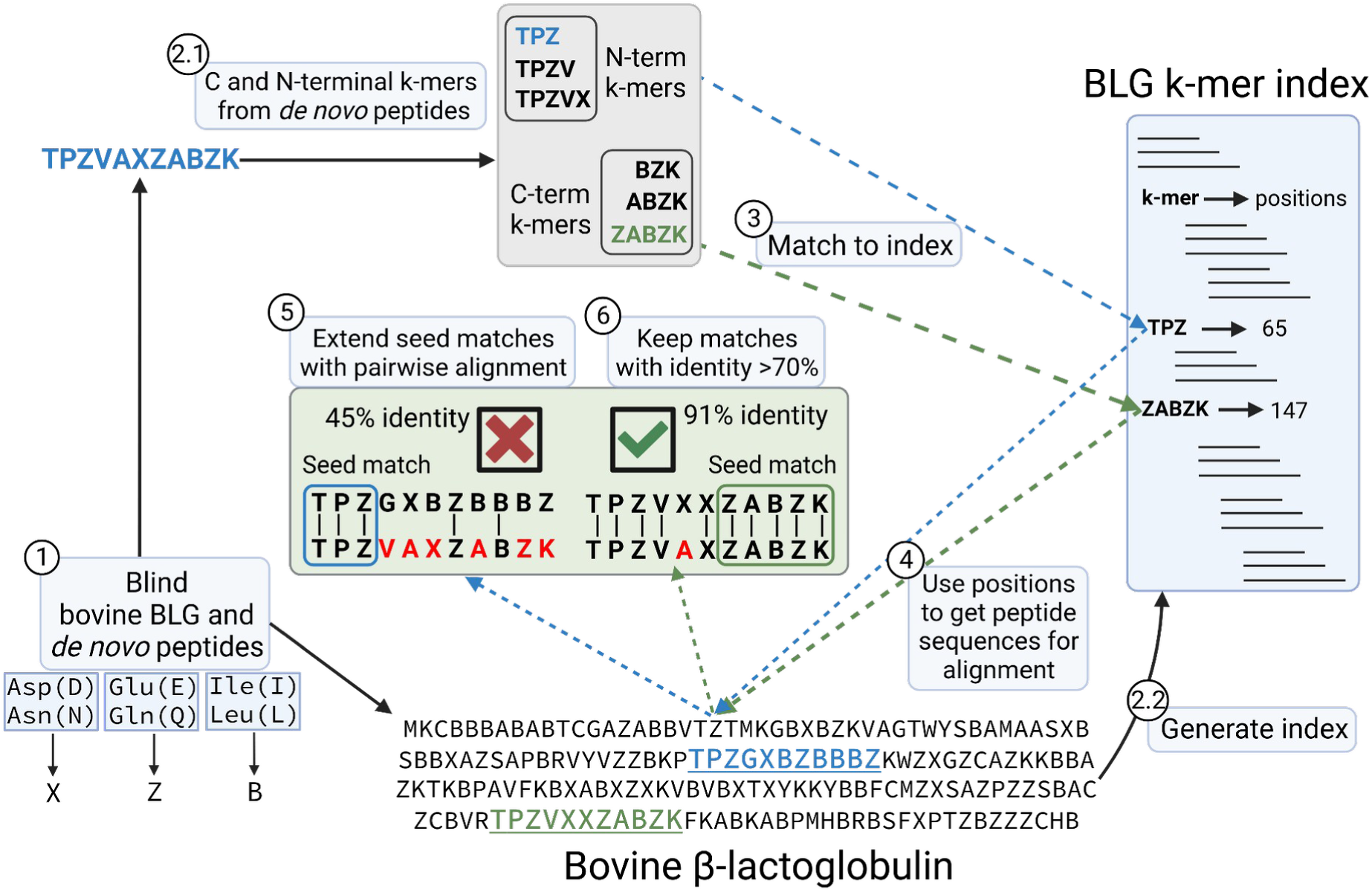
Schematic of the procedure to map the *de novo* peptides to the bovine β-lactoglobulin sequence, while allowing sequencing errors. 1) *de novo* peptides and BLG are ‘blinded’. 2.1) C and N-terminal k-mers are extracted from the peptides. 2.2) BLG index is generated, with k-mers as keys pointing at the positions in the protein. Positions are with respect to the immature BLG containing the signal peptide 3) In this example, the peptide TPZVAZABZK matches 2 different k-mers from the BLG k-mer index, which come from positions 65 and 147 of the BLG product sequence. 4), 5) This results in two possible alignments of the query peptide against BLG. 6) The alignment with the highest accuracy is kept and overall, only alignments with identity above 70% are considered. In case of ties, the alignment with the longest seed match and coming from a C-terminal k-mer would be kept.

### Peptide physico-chemical properties

We extracted or calculated several physico-chemical properties of the BLG: B-factors, relative solvent accessibility (RSA), amyloid propensity and isoelectric point.

We extracted the B-factors from the crystallographic structure of the BLG from PDB accession 7KOT. When determining the three-dimensional structure of a protein using X-ray crystallography, the B-factors are calculated from the attenuation of X-ray scattering due to thermal displacement of the atoms (Sun et al., 2019). B-factors have been extensively used to measure the flexibility of the backbone and side chains of proteins (Sun et al., 2019). The solvent accessible surface area (ASA), in Å^2^, was extracted from the 7KOT structure from PDB using DSSP (Kabsch & Sander, 1983; Joosten et al., 2011). The RSA was calculated by dividing the ASA by the MaxASA obtained from (Tien et al., 2013). We calculated the isoelectric point of all identified peptides using the *pI* function in Pyteomics. For reference, we also calculated the isoelectric point along the BLG sequence using a sliding window of size 12. To estimate the probability of amyloid formation, we used 3 different software, APPNN (Família et al., 2015), AmyloGram (Burdukiewicz et al., 2017) and ANuPP (Prabakaran et al., 2021).

## Results and discussion

The LC-MSMS produced a total of 8,731 MS2 spectra for the 0 days sample and 9,108 and 10,141 for the 4 and 128 days respectively. The increase in the total number of spectra is consistent with the higher non-enzymatic hydrolysis expected in the more degraded samples, which would yield a wider variety of peptides.

### Is FDR enough?

We calculated q-values for each PSMs in each database search software run and sample using pyteomics *q_value* function. Then we calculated the fraction of PSMs above a range of q-values from 1·10^-5^ to 5·10^-2^ and plotted the fraction of identified spectra (Figure 3). Supplementary Table S1 summarises the absolute number of PSMs for each run and software at FDR 0.01 and 0.05, which are highlighted as a blue dashed line in Figure 3, while Supplementary Figure S1 shows the same information only for FDR 0.01. We observe several trends:

1. Looking at each software separately, the fraction of identifications above the same FDR threshold, first slightly increases in the 4-days sample but then decreases in the 128-days.
2. Within the same software, on the 0 days sample, tryptic or semi-tryptic searches give the highest number of identifications, and only as degradation increases, in the 4 and 128 days, semi-tryptic, and then non-specific searches identify more spectra.
3. In each software and sample, the sets of parameters producing more identifications are always the ones using the smaller, dairy-specific DB1.
4. At the least stringent FDR, 0.05, and in the best case, less than 60% of the spectra is identified for the 0 and 4 days samples. For the 128 days sample, this is reduced to less than 30%.

**Figure 3.**
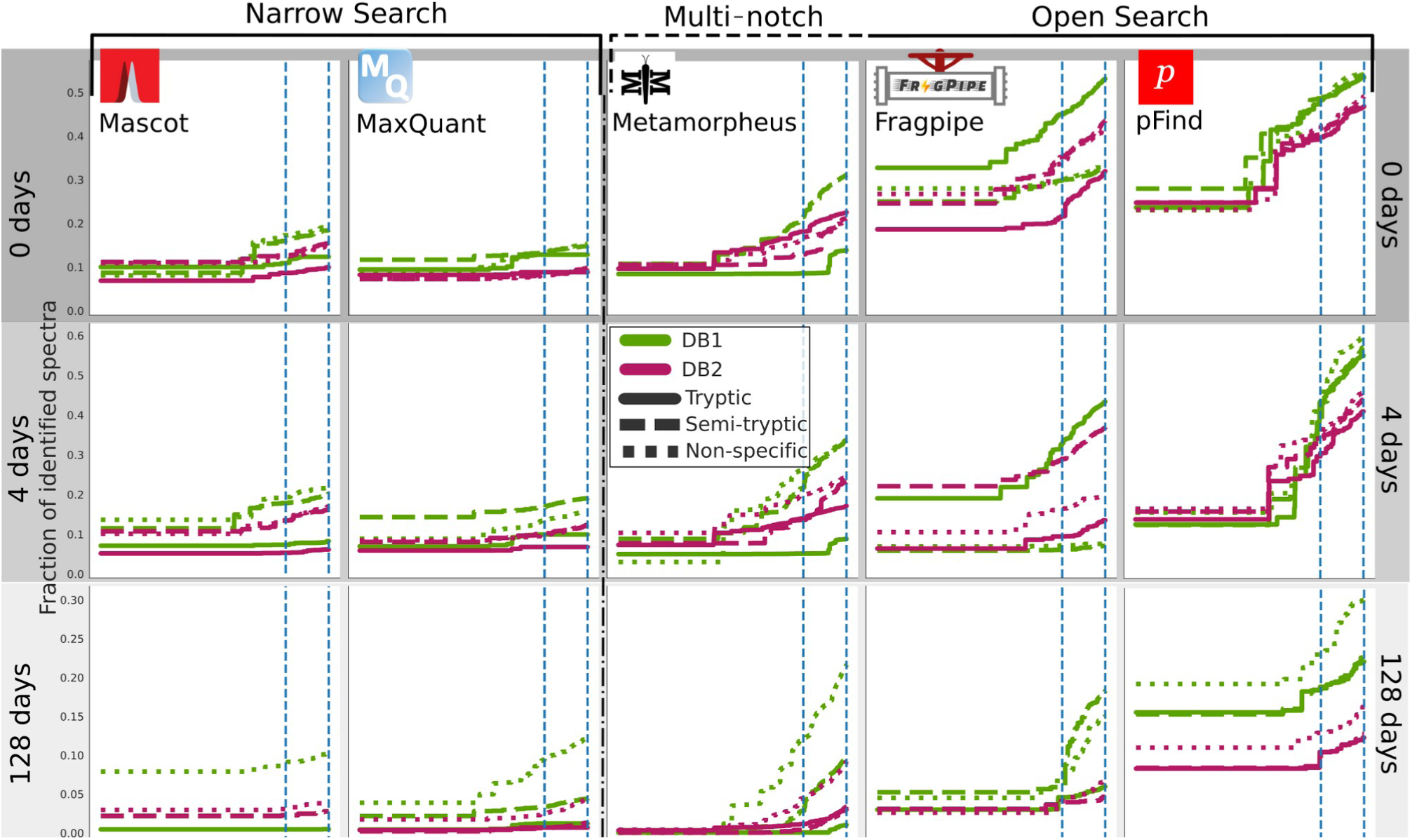
Fraction of identified spectra for FDR thresholds from 0 to 0.05 (at 10^-5^ step) for *DBs* software. The rows represent the increasing heating time (0, 4, 128 days), with each software represented in a column: MaxQuant and Mascot are narrow search software, Metamorpheus multi-notch and Fragpipe and pFind open search. Dairy database searches (DB1) searches are represented in green, and bovine proteome searches (DB2) in magenta. The solid lines are for tryptic searches, dashed for semi-tryptic and dotted for non-specific. The blue dashed lines mark the 0.05 and 0.01 FDR thresholds that are usually applied.

The amount of identified PSMs at a given FDR are widely affected by the search space (Figure 3), controlled by the choice of the database and enzymatic search. And in general, it is remarkable that for such a simple system explored in this study, vast amounts of spectra remain unidentified. This is more pronounced for the most degraded and therefore complex sample (Figure 3). MaxQuant and Mascot identify less than 20% of spectra for the 0 days and 10% for the 128 days. This is significantly lower than the other software, almost for any parameter set. This makes them particularly affected by the decrease in the amount of PSMs when using the larger database.

The main reason behind the reduction on the number of PSMs when we expand the search space is already known in proteomics and in statistics general: if by Increasing the search space, the proportion of peptides in the database that are not present in the samples also increases, there will be more chances of random and decoy matches with high scores (Jeong et al., 2012; Noble, 2015). Therefore, to achieve the same desired FDR, the score threshold to accept positive matches becomes more stringent, rendering fewer accepted PSMs (Jeong et al., 2012; Noble, 2015). This means that large databases containing a large number of proteins that are not present in the samples, like DB2, inadvertently boost the FDR and apparently reduce the number of identifications. Something similar applies to the use of non-specific searches for samples that are well-preserved and less degraded. This adds to the inherent divergence between FDR and FDP of the TDC (He et al., 2015; Couté et al., 2020; Madej & Lam, 2023; Ebadi et al., 2023). But the answer to this problem cannot just be to use small, targeted databases, even though it would reduce the computing time and resources and might produce more PSMs. We can speculate what would happen in the opposite scenario, using a very small database against a complex sample, for example dental calculus. To start, we might miss some of the contents of the sample. Another more inadvertent effect is that wrong, low-quality matches between a spectrum and a peptide might surface with a low enough FDR (since the search space is small). This is further pronounced if the correct peptide, which could have competed and achieved a better PSM, is absent in the database. This is why common contaminants are included in the searches, even though we are not interested in them.

In general, we want the set of proteins and peptides that are in a sample and the set of what we are looking for (search space) to match as closely as possible (Jeong et al., 2012; Noble, 2015). While the content of a sample is not known *a priori*, it is clear in our simple system that using the whole bovine proteome to interrogate our data is utterly unnecessary and counter-productive. Going into a more realistic scenario, with real archaeological samples, having a clear research question and hypotheses and a good understanding of the historical, archaeological, diagenetic and taphonomic context will always help to delineate the optimal database and parameters given the compromises exposed. Studies like those of Wilkin et al. (2020, 2021), Bleasdale et al. (2021) and Hendy et al. (2018; 2018) are examples of different strategies and challenges regarding the complexity of samples and scope of the databases. They highlight the difficulty of building an unbiased and targeted database that encompasses all the contents of a sample. Targeted databases for different types of tissues like bone or enamel, microbiomes or other mixtures like foodcrusts need to be further developed.

Finally, another mechanism that reduces the amount of PSMs is the exclusion of correct matches because of a poor separation of the score distributions between the target and decoy matches. The low amount of identifications at any FDR on MaxQuant and Mascot, and the dramatic change when switching from 0.01 to 0.05 on Fragpipe, Metamorpheus, and pFind to a lesser extent (Figure 3 and Supplementary Table S1), indicates this is likely happening here. The choice of a score metric for the PSMs is of great importance, because a good scoring function will be able to assign low scores to wrong matches, and vice versa. Therefore, wrong matches can be excluded without eliminating correct matches. Furthermore, a subsequent classifier can be trained to separate the score distributions of the target and decoy hits, integrating different metrics such as the primary search score, peptide length, misscleavages, or PTMs. The use of post-processing classifiers is advised to set up accordingly and is now routinely integrated in most software pipelines: Fragpipe uses PeptideProphet (Keller et al., 2002; da Veiga Leprevost et al., 2020) or Percolator (The et al., 2016) (not used in this study, as it is not compatible with open searches), pFind integrates a linear classifier (Chi, Liu, Yang, Zeng, Wu, Zhou, Wang, et al., 2018). MaxQuant uses Posterior Error Probabilities (PEP) in a similar way (Tyanova et al., 2016) and Mascot also integrates Percolator in its pipeline.

### Open search vs Narrow search

In the previous section, we commented on the implications of the database and enzyme digestion choices in the general TDC approach and FDR. However, the presence of a wide variety of PTMs in ancient, degraded proteins, offers further challenges to identification.

The largest differences in the number of PSMs are between Narrow and Open search software (Figure 3 and Supplementary Figure S1 Table S). Open search furnishes more identifications in all 3 samples, provided that DB1 is used with the appropriate enzymatic digestion (or lack). MaxQuant, Mascot and Fragpipe-Narrow tend to have a similar number of PSMs across the samples and enzymatic cleavage and behave similarly in terms of which settings produce more PSMs for each sample. On the other hand, the open search software has more variability, especially open-Fragpipe (Figure 3, Supplementary Figure S1 and Supplementary Table S1).

Open search is capable of identifying spectra from peptides bearing PTMs that are not accounted for and therefore remain unidentified (Chick et al., 2015). This is so because, in practice, the search space in this software is more unconstrained, while at the same time implementing algorithms that allow a fast search. FragPipe-open finds mass shifts not corresponding to the variable or fixed PTMs set, in 46.6%, 31.8% and 17% of the PSMs below FDR 0.05 for the 0, 4 and 128 days, respectively. The decrease in the identification of PTMs with the more degraded samples might seem counterintuitive. Nonetheless, the backbone cleavage and accumulation of PTMs in the 128 days samples might be so extensive that the Open search cannot even identify those spectra, and only a smaller fraction of the modified peptides is identified. In addition to the problems with FDR already commented, Kong et al. (2017) reported inherent problems with the TDC approach when using narrow search and an increase in the statistical power of open searches. They also reported that 68% of spectra reassigned by open search MSFragger to a different peptide from narrow search, had greater support in the former. They suggest there are potential false positives in spectra matched to unmodified peptides. However, Freeston et al. (2024) explored the trade-offs and discrepancies between open and narrow searches and dispute these claims. They attribute this to the presence of chimeric peptides and flaws in FDR control in PeptideProphet and Percolator. They found that narrow searches can perform better at identifying unmodified peptides. Fragpipe’s performance with respect to its narrow search counterpart is lower in non-specific search of the 128 days sample and non-specific and semi-tryptic for the 4 days sample. This, as well as Fragpipe’s more unstable performance, might be a consequence of the findings by Freeston et al. (2024). Like Fragpipe, pFind also includes a rescoring or reranking step, while Metamorpheus only relies on the primary search score to discriminate between true and false matches (Wenger & Coon, 2013).

In order to highlight the differences between algorithms within Open search, we created UpSet diagrams in Figure 4A with the common spectra identified in the different open search software and enzyme cleavage runs and for each of the samples. For FragPipe and Metamorpheus, on the 128 days sample, most spectra are identified only on the non-specific search, and the difference between the 3 types of digestion increases with sample degradation (Figure 4A). In contrast, in pFind most identified spectra are common in the 3 searches throughout the 3 samples, although the totals decrease (Figure 4A). Moreover, most of the spectra identified by Metamorpheus and Fragpipe are also identified by pFind, while few spectra are exclusively identified by Fragpipe and/or Metamorpheus. Open-pFind takes a different approach than MSFragger or Metamorpheus; it finds k-mer tags in the spectra that are extended until there is a match to the peptide database, and given a mass shift (Chi, Liu, Yang, Zeng, Wu, Zhou, Niu, et al., 2018). Open-pFind is capable of finding non-tryptic peptides even in tryptic search in the 128 days sample (Figure 4A). The mass shift between the precursor ion and the peptide candidate is treated as a potential modification, and PTMs are proposed by testing all valid positions according to Unimod (Creasy & Cottrell, 2004).

**Figure 4.**
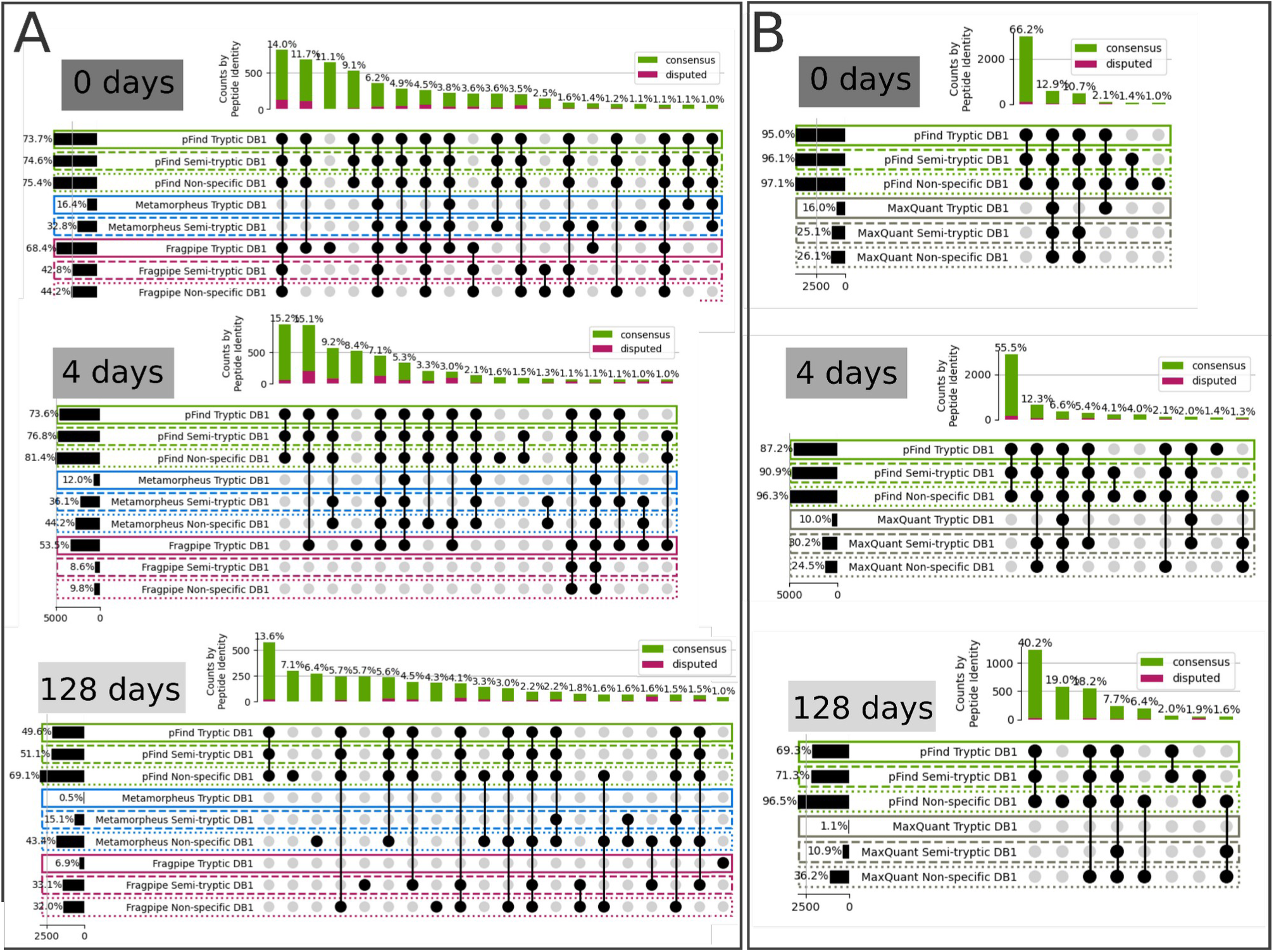
UpSet plots by sample showing common identified PSMs across software and enzymatic cleavage. PSMs below FDR 0.05 are considered. A) UpSet plots for pFind (green), Metamorpheus (blue) and Fragpipe (pink) and tryptic (solid line), semi-tryptic (dashed) and non-specific (dotted) searches. B) MaxQuant and pFind UpSet plots. All cases are with the dairy database (DB1). The vertical bars indicate the sizes of each set and the percentages are with respect to the size of the union of all sets. The degree of agreement between the peptide sequences of each set is represented in red (average pairwise Levenshtein distance less or equal than 1) and green (average pairwise Levenshtein distance greater than 1).

In Figure 4B we compared pFind and MaxQuant using UpSet diagrams, again considering PSMs below 0.05 FDR. The sets of spectra identified by MaxQuant are always contained in the set identified by at least one pFind run. In the most extreme case, for the 128 days sample, only 8% of the spectra are identified by the semi-tryptic or non-specific runs in MaxQuant, and not by the tryptic run on pFind (Figure 4B).

Open-pFind, even in the tryptic search on the most degraded sample, “unlocks” a spectra space that the other software does not identify. And in general, the Open search software provides identifications for spectra for which narrow search does not. We found that those PSMs exclusively identified by pFind tend to have a slightly lower score than those also identified by MaxQuant (Wilcoxon rank-sum test *p-value=0*). But even though significant, the magnitude of the difference remains small and many pFind-exclusive PSMs score high (Figure 5).

**Figure 5.**
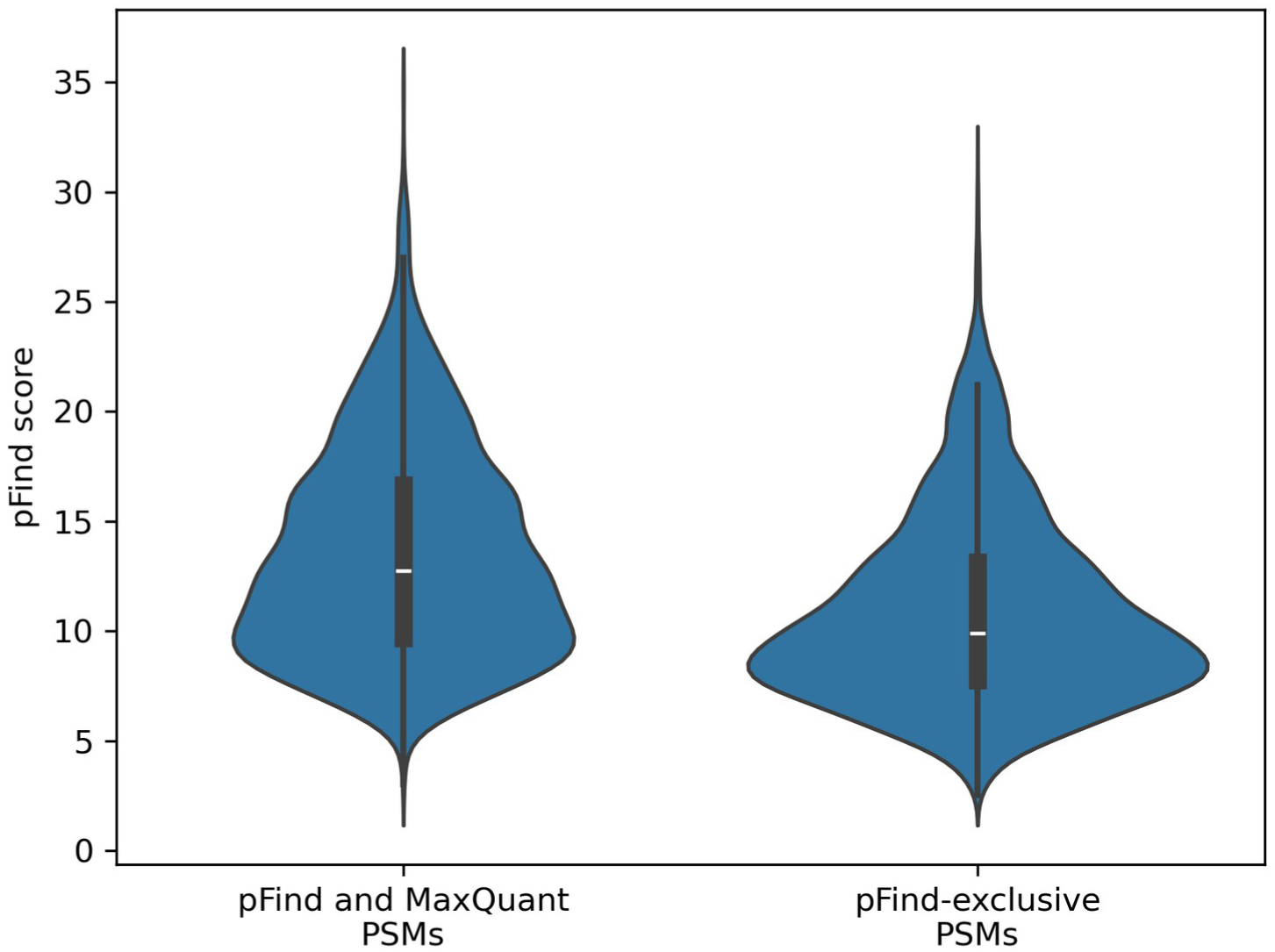
Distribution of the pFind scores for PSMs below 0.05 FDR identified both by pFind and MaxQuant and only identified by pFind.

For the cases, when they both provide a peptide identification for the same spectra, we explored to what extent the pFind and MaxQuant agree. Overall, the level of agreement decreases on the 128-days sample when running semi-tryptic or non-specific searches (Figure 6, left column bar plots). This is, with the more complex sample and unconstrained search space. We also compared the Andromeda and pFind scores for each PSM and the Levenshtein distances (the extent of disparity) between the identified sequences (Figure 6, right column scatter plots). There is an overall correlation between the scores, with those having the highest Levenshtein distances showing low degree of correlation, or scoring low in one or both software (Figure 6). However, disputed peptide sequences still overlap broadly with those where there is a consensus.

Lastly, we compare the running times between pFind, FragPipe and MaxQuant. It has already been reported that the 2 open search software run much faster than other narrow search ones (Kong et al., 2017; Chi, Liu, Yang, Zeng, Wu, Zhou, Niu, et al., 2018). Here, we also run a subset of the MaxQuant runs on the Cambridge HPC (Table 1) with the same amount of CPUs in order to compare with Fragpipe in the same system. MaxQuant and pFind were executed on the same system (Table 1). We found that pFind runs about 40 times faster than MaxQuant on MiniMax and FragPipe about 30 times on Cambridge HPC.

**Figure 5.**
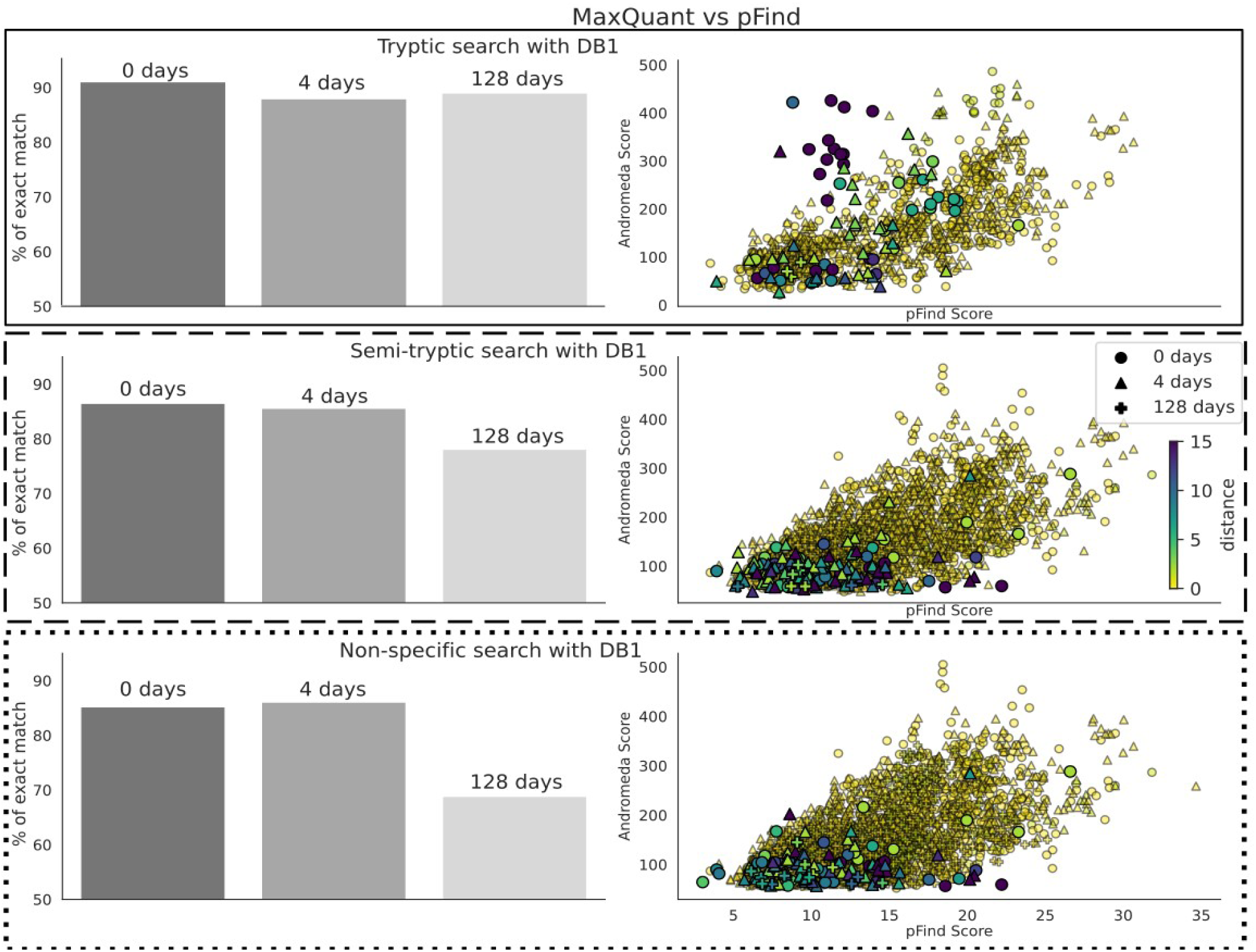
Comparison of the identifications made by Fragpipe and MaxQuant using the dairy database and tryptic search (top panels, with a solid outline), semi-tryptic (middle, with dashed outline) and non-specific (bottom with dotted outline). When both software identify the same spectra, we calculate the distance between the peptide sequences, which is expected to be 0. Panels on the left are a histogram of the Levenshtein distances, split by sample (0, 4 and 128 days). The text is the percentage of common spectra for which both sequences are identical. Panels on the right are the scores given by the software to that same PSM. The color of each point is again the Levenshtein distance.

The abundance and range of PTMs is likely to be limited in this study, given the low complexity of the sample. For example, there is no lactose that would generate lactosylations (Czerwenka et al., 2006; Bosman et al., 2021). However, on a real, complex, archaeological or palaeontological ancient sample, we can expect a plethora of post-translational modifications, from the ones occurring *in vivo,* to those induced by human activities, bacteria and time. A simple narrow search, as seen here and as frequently reported in the literature and by researchers, leaves massive, and potentially very informative, amounts of data unexplored. As we will comment in further detail, we recommend including open search software in the analysis workflow in Palaeoproteomics; at least for PTM discovery to augment narrow searches, similarly to what Metamorpheus G-PTM-D does (Solntsev et al., 2018).

### *De novo sequencing* coverage bias and accuracy

We have seen that when we use external knowledge and data to search the spectra, we are always required to make choices that will affect the number, quality and significance of the identifications. Open search software can search for unspecified PTMs. But *de novo* sequencing (*dnS*) software can in theory reconstruct the peptide sequence just from the spectra without the use of an external database. However, sequencing errors might arise (Muth & Renard, 2018; Beslic et al., 2023), which, without external validating data, can have a high impact in the discovery of potential novel peptides from extinct or unsequenced organisms and phylogenetic studies. In this section we look at the interplay between degradation, sequence coverage bias and accuracy of *dnS*. We also discuss how the choice of the database affects the accuracy of the results in this case, since DirecTag and PepNovo+ rely on it.

Since we have a very simple system, with a single and known protein, it is easy to map the *de novo* sequenced peptides back to the BLG to obtain sequence coverage and also identify sequence errors. We follow the procedure explained in <u>Mapping of de novo peptides</u> and Figure 2. Finally, peptides with less than 70% overall identity with the BLG sequence are discarded. We found that overall, alignments coming from C-terminal seed matches have a greater identity than those from N-terminal k-mers, when none of them are given priority (Figure S3). This is in line with previous studies reporting that long *b-ions* are unstable and therefore *y-ions* are preferentially generated in HCD (Shao et al., 2014; Medzihradszky & Chalkley, 2015). Therefore, HCD produces better ion fragment coverage on the C-terminal and in turn less sequencing errors.

As with *DBs*, we studied the effect of the size of the sequence database for *de novo* methods that use it at any point of the pipeline, like DirecTag and PepNovo+. When the larger DB2 is used in DeNovoGUI, DirecTag and PepNovo+ attempt at covering larger parts of the BLG sequence, but at the cost of decreasing the overall accuracy of the peptides (Figure 7A, B, C and Supplementary Figure S2). In contrast, Novor achieves almost identical results (Figure 7A and Supplementary Figure S2). Within DeNovoGUI, DirecTag and PepNovo+ rely on an external database, either for a tagging hybrid method or for a subsequent MS-BLAST, respectively. We can draw a parallel observation as when the FDR is inadvertently boosted in *DBs* if the database contains many irrelevant sequences. In this case, incorrectly *de novo* sequenced peptides or tags are queried against a plethora of sequences from DB2, resulting in spurious matches that end up validating or wrongly extending them. Novor does not use a database at any point, but the overall amino acid accuracy is always lower (Figure 7A and Supplementary Figure S2). Beslic et al. (2023) also reported Novor performed worse than other *dnS* software in terms of amino acid and peptide recall.

**Figure 7.**
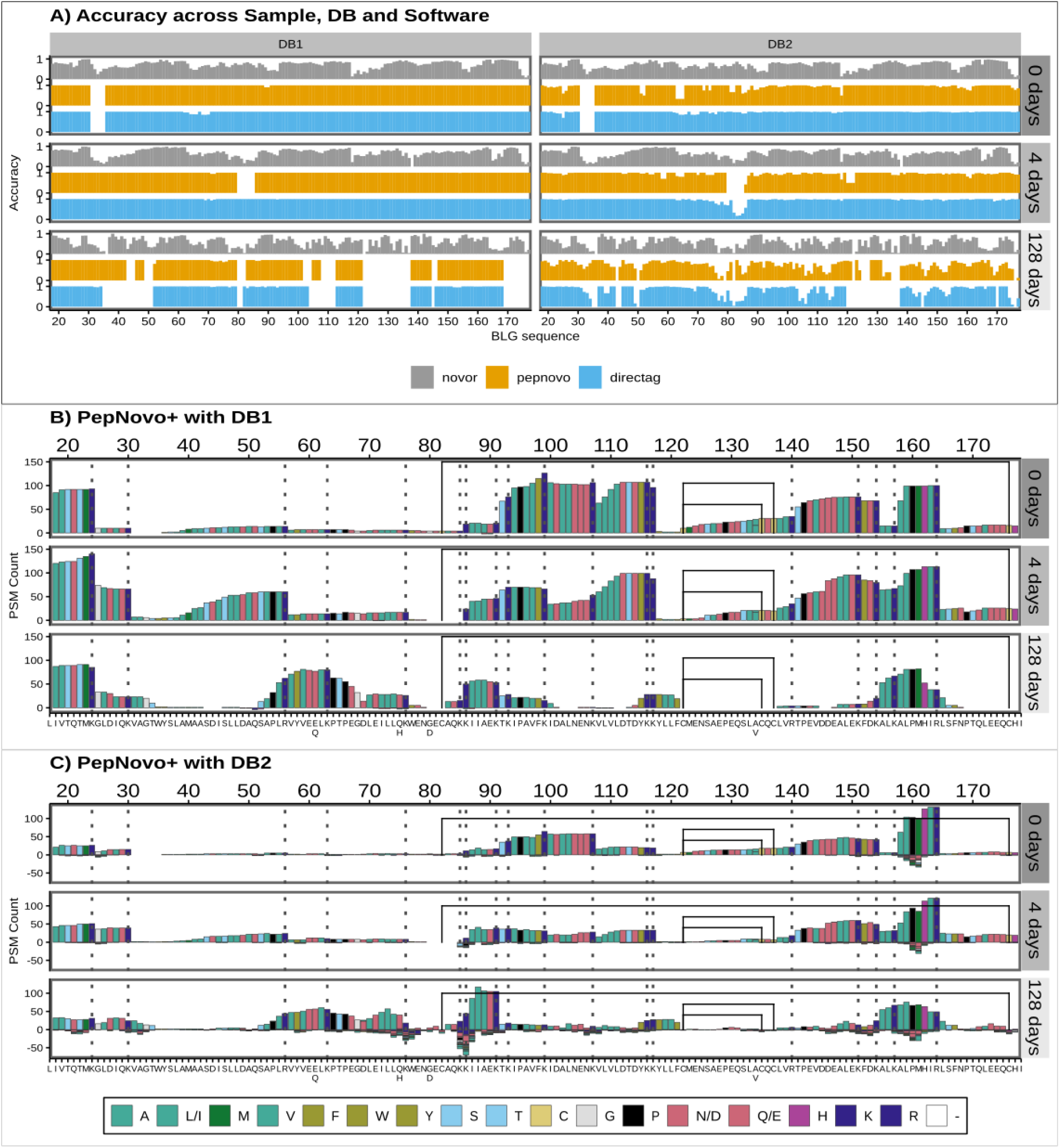
Positional sequence coverage and amino acid accuracy. Coverage is measured in the y-axes as the number of times an amino acid position is covered by an identified peptide, while accuracy is the proportion of those that are correctly identified. The BLG signal peptide is not shown, but positions are given counting it, and therefore start at 17, just to highlight that it would not be present in the mature BLG found in milk. Panel A) shows the amino acid accuracy for the 3 *de novo* tools using either the dairy database (DB1) or the bovine proteome (DB2). Panels B and C show the coverage for PepNovo+ using DB1 and DB2 databases. Values below 0 correspond to mismatches between the peptide sequence and the actual BLG sequence. In panels A and B, the grey dotted lines mark the tryptic sites and the black lines are the disulfide bonds between Cysteines in positions 82 and 176, 122 and 135 and the alternative 137. Amino acids are coloured by main properties: hydrophobic ones (A, L/I, M, V) are in teal green, with an olive shade for the aromatic ones (F, W; Y). Acidic amino acids (D and E) are in rose (or their amide group counterparts N and D, as they are not differentiated here), while basic ones are in dark blue (K and R) or purple (H). Other hydrophilic amino acids (S and T) are in cyan. Cysteines (C) are highlighted in yellow, glycine (G) in grey and proline (P) black. Colour scheme source: https://personal.sron.nl/~pault/

To study sequence accuracy along the BLG we take every BLG-assigned sequenced peptide position and 1) count how many times it is observed and 2) identify if it is correct or not in order to calculate the accuracy at each position. Incorrect amino acids produce a negative count (Figure 7 B, C). For Novor, inaccuracies accumulate with the more degraded samples (Figure 7A). In contrast, DirecTag and PepNovo+ decrease their coverage, but inaccuracies only occur when using the DB2. These inaccuracies are not spread evenly across the BLG, but are more pronounced in specific areas (Figure 7A, B, C). Moreover, regions presenting good coverage, between positions 80 and 90 or around 160, also present the highest number of wrong observations (Figure 7 C)

In line with the database effect, DirecTag and PepNovo+ are more conservative and overall present less hits (150 for the best position), because sequencing errors are prevented and wrong peptides discarded upon matching against a database. Novor, in contrast, attempts at sequencing everything and has improved coverage but returns more erroneous peptides.

*De novo* sequencing is a critical tool in palaeoproteomic studies looking at the phylogenetic relations of extinct animals (Welker et al., 2015, 2019; Chen et al., 2019; Presslee et al., 2019; Cappellini et al., 2019). Crucially, these studies use PEAKS, MaxQuant and BLAST, in sequential workflows tailored to the identification and validation of single amino acid polymorphisms and novel peptides. The combination of hybrid methods and *dnS* and *DBs* reflects the compromise between the discovery of novel sequences and the minimization of sequencing errors. As seen in Figure 7, sequencing errors can accumulate in highly degraded proteins and regions. Software that does not rely on any external database is more affected by this. At the same time, a database can be a curse, rather than a blessing, if it contains too many irrelevant sequences, like DB2. Therefore, these studies use databases restricted to a specific species, or type of protein like collagen or tissue, like bone or enamel proteomes.

### Peptide chemical properties and coverage bias

In the previous section, we reported a coverage bias for *dnS*, which also changes as degradation proceeds. Figure 8 shows the coverage for pFind tryptic search (A) and MaxQuant with tryptic (B) and semi-tryptic (C) search with the dairy specific DB1. There are sections of the sequence and peptides with a much greater support than others, meaning they were identified in more different PSMs. In the 128 days sample, this coverage bias dramatically changes and some regions become “invisible”, while others increase their relative coverage. In line with the findings of previous sections, pFind tryptic search and MaxQuant non-specific search coverage of the 128-days sample is similar (Figure 8A and C), while MaxQuant’s tryptic search fails to identify broad sections of the BLG (Figure 8B). We explored whether structural flexibility, relative solvent accessibility (RSA), amyloidogenicity and isoelectric point along BLG protein sequence and 3D structure correlate with these patterns. For this we extracted the B-factors and relative solvent accessibility from the 7KOT 3D structure from PDB. We estimated the amyloid formation propensity using different algorithms and predictors and calculated the isoelectric point (Figure 9).

**Figure 8.**
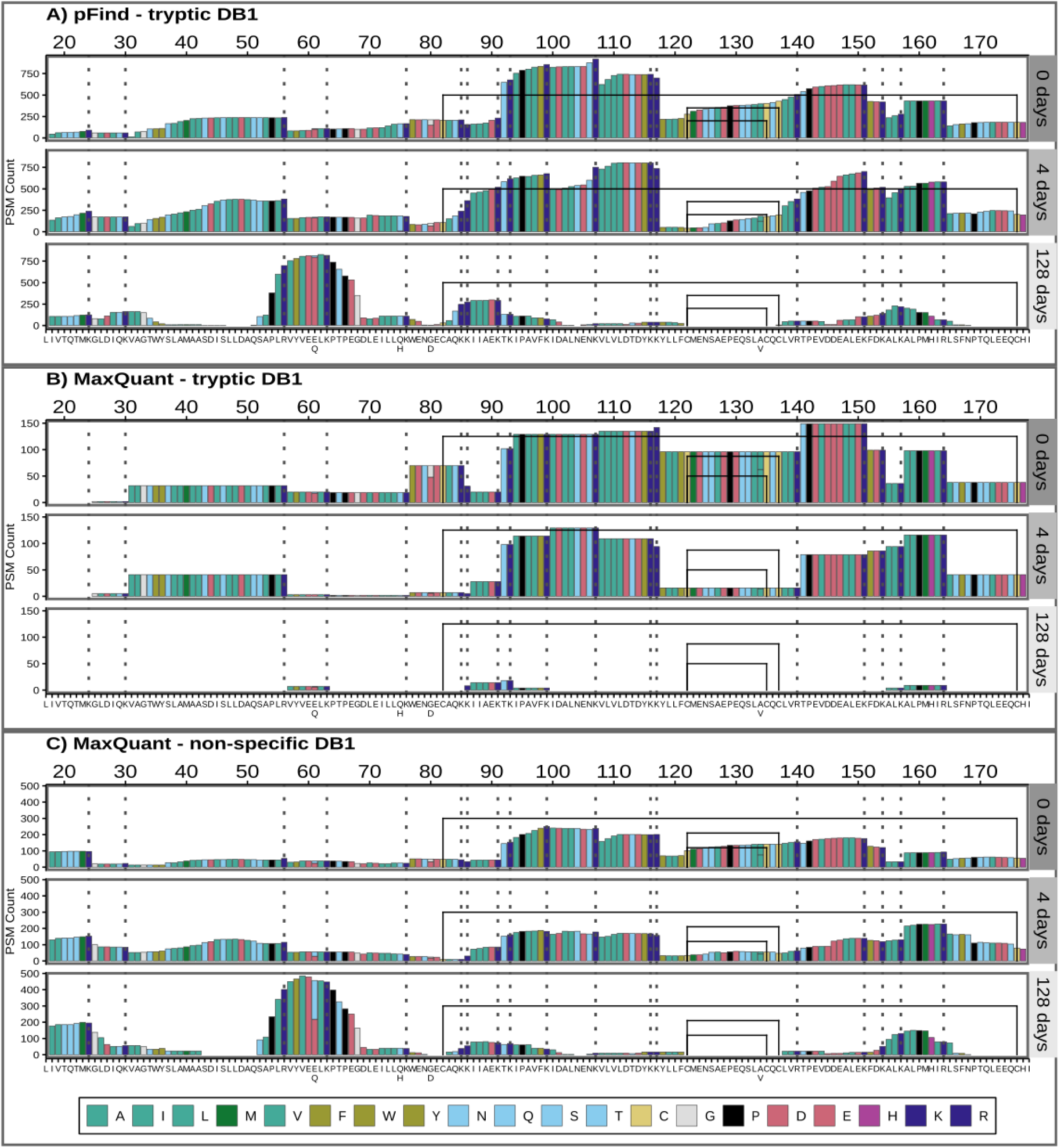
Sequence coverage for pFind tryptic search (A) and MaxQuant with tryptic and non-specific search (B and C). The BLG signal peptide is not shown, but positions are given counting it, and therefore start at 17, just to highlight that it would not be present in the mature BLG found in milk. PSMs below 0.05 FDR are included. Amino acids are coloured by main properties: in teal green hydrophobic ones (A, I, L, M, V), with an olive shade for the aromatic ones (F, W, Y). Acidic amino acids (D and E) are in rose, while basic ones are in dark blue (K and R) or purple (H). Other hydrophilic amino acids (N, Q, S and T) are in cyan. Cysteines (C) are highlighted in yellow, glycine (G) in grey and proline (P) black. Colour scheme source: https://personal.sron.nl/~pault/

The region between positions 85 and 117 has the greatest coverage in the non-degraded (0 days) and moderately degraded (4 days) samples. It coincides with a region with a high isoelectric point, several exposed tryptic sites and low probability of amyloid aggregate formation. However, after extensive degradation its coverage greatly decreases and some highly flexible sections (97 to 117) are barely identified. In contrast, the relative coverage of the region between positions 53 and 67 increases only after extensive degradation and using non-specific search. The only 2 tryptic sites here are relatively not as exposed and the isoelectric point is low.

The region between positions 105 and 140, features compact and rigid beta-sheets connected by flexible loops in the 3D structure, with high propensity of forming amyloid structures, it is negatively charged (low isoelectric point) and lacks tryptic sites (Figure 9). This region’s coverage is overall reduced from the 0 to 4 days samples and then is not observed in the 128 days sample.

**Figure 9.**
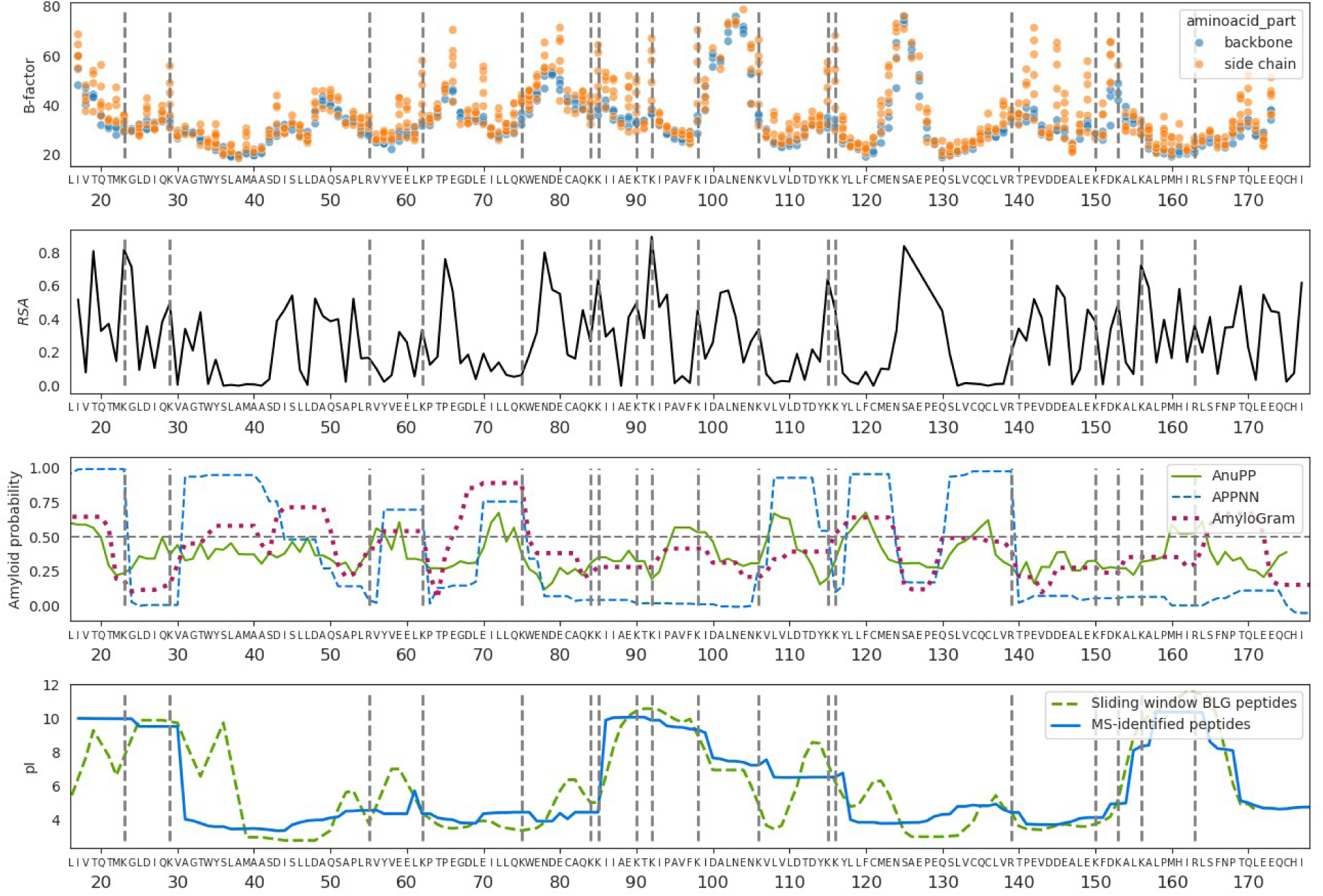
Physico-chemical properties along the beta-lactoglobulin. The top graph are the B-factors of all atoms except hydrogens. The carboxylic and alpha carbons and nitrogens from the amino group along the backbone are in blue, while other side chain atoms are in orange. The second graph shows the relative solvent accessibility of the BLG folded structure. The third panel is the positional amyloid formation propensity estimated by AnuPP (Prabakaran et al., 2021), ANNPP (Família et al., 2015) and AmyloGram (Burdukiewicz et al., 2017) software. The last graph is the isoelectric point. The tryptic sites (lysines and arginines) are marked as grey dashed horizontal lines.

It is therefore a combination of processes and mechanisms that will determine whether a protein or peptide will be successfully extracted, detected in the mass spectrometer and identified (Evans et al., 2024). First there needs to be well exposed tryptic sites (if using trypsin and tryptic or semi-tryptic search), otherwise, those regions can only appear after denaturalisation and hydrolysis of the backbone (128-days sample) (Evans et al., 2024), and then only observed using semitryptic or non-specific search. However, peptides can start forming aggregates after they are enzymatically cleaved or hydrolysed (4 days), and highly flexible, polar and charged regions are also more vulnerable to being hydrolyzed by water (Fonseca et al., 2024). At the same time, other properties might protect peptides from hydrolysis and degradation. The peptide T_141_PEVDDEALEK_151_ is one of the most encountered BLG peptides in the literature (Wilkin et al., 2020, 2021); (Bleasdale et al., 2021). It presents highly flexible and negatively charged side-chains (Figure 9). Here, in contrast, it is hardly identified after extensive degradation. This suggests that the real conditions in which the peptide is found, mainly the presence of a mineral surface, contributes to its preservation, as it has been suggested using molecular simulations (Hagiwara et al., 2009; Fonseca et al., 2022). The mineral surface and other characteristics of a real sample also determines the extent to which proteins are successfully extracted in the laboratory, which is not discussed here.

The last main factor determining the detection of peptides is the ionisation efficiency by the electrospray. Peptides with high isoelectric point are already positively charged and therefore easier to be detected when running in the electrospray in positive ion mode. Negatively charged peptides, with low pI, will depend entirely on ionisation efficiencies at the electrospray (Liigand et al., 2017).

### Towards an integrated Paleoproteomics pipeline

Results can vary greatly between MS/MS identification strategies, even for such a simple system as the one explored here. Moreover, there are challenges that each method cannot single-handedly solve. To tackle this, we devise an standardised approach that combines the powers of the 3 methods discussed (*de novo,* open and narrow search), suited for the study of ancient proteins. This has already been done in other more informal and *ad-hoc* approaches that combine *de novo* sequencing and database search (Welker et al., 2015, 2019; Chen et al., 2019; Presslee et al., 2019; Cappellini et al., 2019). Meta-search engines like DeNovoGUI (Muth et al., 2014), SearchGUI (Barsnes & Vaudel, 2018) or platforms like PeptideShaker (Vaudel et al., 2015) focus on aggregating results from different software. This is important to catch disputed sequences or give more reliability to the consensus ones.

The pipeline would integrate the sequence discovery of *de novo* sequencing with the PTM discovery of open search, and a process to narrow down sequence databases. These can be used to (1) refine a protein sequence database and (2) expand the PTMs to search at a final narrow search (Figure 10) that would be more reliable. At the end, peptide identifications by different software can be integrated to increase the confidence of the results.

**Figure 10.**
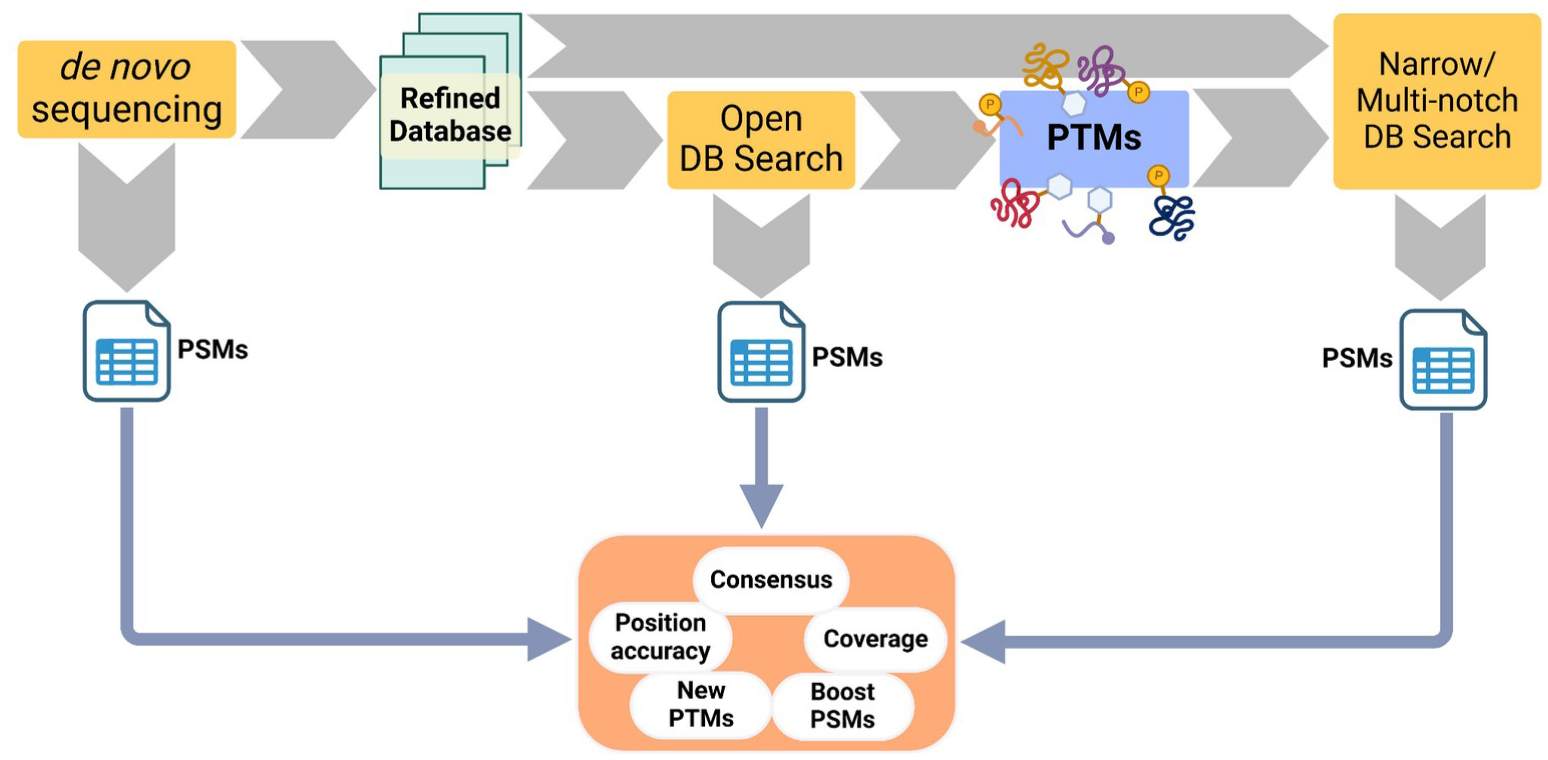
Proposed pipeline for the analysis of ancient proteins. It starts with a *de novo* sequencing step that refines or produces a targeted database. This database is then used for an open search and the novel, abundant or relevant PTMs are extracted. The refined database and the PTMs are then fed into a closed or a multi-notch database search. At the end the PSMs of the 3 software runs are combined, to check for consensus or disputed sequences, study position accuracy and coverage. This would increase the confidence of the results, while producing more PSMs than each of the steps alone, and identify PTMs reliably.

Moreover, we believe it is useful to study sequence coverage and positional information to identify commonly identified regions and inaccuracies. Finally, the identification of a wider array of PTMs can enrich the information that we can gather from ancient proteins in archaeological and palaeontological samples.

### Conclusion

Palaeoproteomics is a maturing discipline that is developing new strategies to overcome its own challenges. Teams working on computational proteomics are constantly releasing new software and updates with new features. These implement a wide variety of underlying methods and algorithms that we broadly group in *de novo* sequencing or database search. Palaeoproteomics frequently borrows methodology and software from the broader field of proteomics. Yet there are unique challenges to applying these tools to palaeoproteomics, such as the difficulties associated with selecting a properly constrained database or accounting and identifying PTMs. It is therefore important to assess the pertinence and validity of the different strategies and parameters in the study of ancient proteins. Here we have done so by using a simple and progressively degraded protein model, β-lactoglobulin, heated at pH 7 for 0, 4 and 128 days.

We highlight the importance of constraining the search space in *DBs* to improve the significance and reduce computing times. Setting the enzymatic digestion according to the level of degradation and using targeted databases are the most important factors. At the same time, Open search software can identify more peptides with PTMs that would have been otherwise unidentified. Moreover, the study of new PTMs, like glycations, can open the door to answering new questions in archaeology. *De novo* sequencing goes one step further in freeing researchers from database selection, which we have shown to hugely impact the results. However, the possibility of sequencing mistakes cannot be disregarded and in practice, external databases and complementary *DBs* are commonly used. Recently, new *dnS* programs like Casanovo (Yilmaz et al., 2022), InstaNovo (Eloff et al., 2023), or π-HelixNovo (Yang et al., 2023) are reporting promising performance. They use transformer neural network architecture, (as used in Large Language Models), which seem suited to protein sequences as they can be seen as another language. A language that is revealed in spectral data, in which the presence or absence of peaks and their intensities are intercorrelated.

As it has already been done in more informal and *ad-hoc* approaches, we propose a standardised approach that combines the powers of the different approaches discussed here. *De novo* sequencing software, especially those using transformers, and Open search can be used to discover novel sequences and PTMs and initially constrain the search space of possible protein contents and modifications in a complex sample. Using this information, a final, constrained *narrow database search* with targeted database and PTMs would boost identifications confidently.

## Supporting information

Supplementary File S1. Parameters

## Data and code availability

Python and R notebooks used to process the *DBs* and *dn*S data can be found in https://github.com/ismaRP/PPbenchmark.git and <u>10.5281/zenodo.13785336.</u> RAW LC-MSMS and PSM data resulting from the analysis with Fragpipe, pFind, Metamorpheus, MaxQuant, Mascot and DeNovoGUI can be accessed in Zenodo <u>10.5281/zenodo.13785293.</u> MaxQuant and Fragpipe parameters and workflow files are also included.

## Acknowledgements

IRP is currently funded by the European Union’s Horizon 2020 Research and Innovation Programme under the Marie Skłodowska-Curie grant agreement No 956410. At the time of producing and writing this work BN was funded by European Union’s Horizon 2020 Research and Innovation Programme under the Marie Skłodowska-Curie grant agreement No. 801199 and with MC by the European Union’s EU Framework Programme for Research and Innovation Horizon 2020 under Grant Agreement No. 787282 (B2C). MC and MM were also supported by the Danish National Research Foundation (DNRF128). YC and JD are funded by the European Union’s Horizon 2020 Research and Innovation Programme under the Marie Skłodowska-Curie grant agreement No. 956351.

## Supplementary Data

**Supplementary Figure S1.**
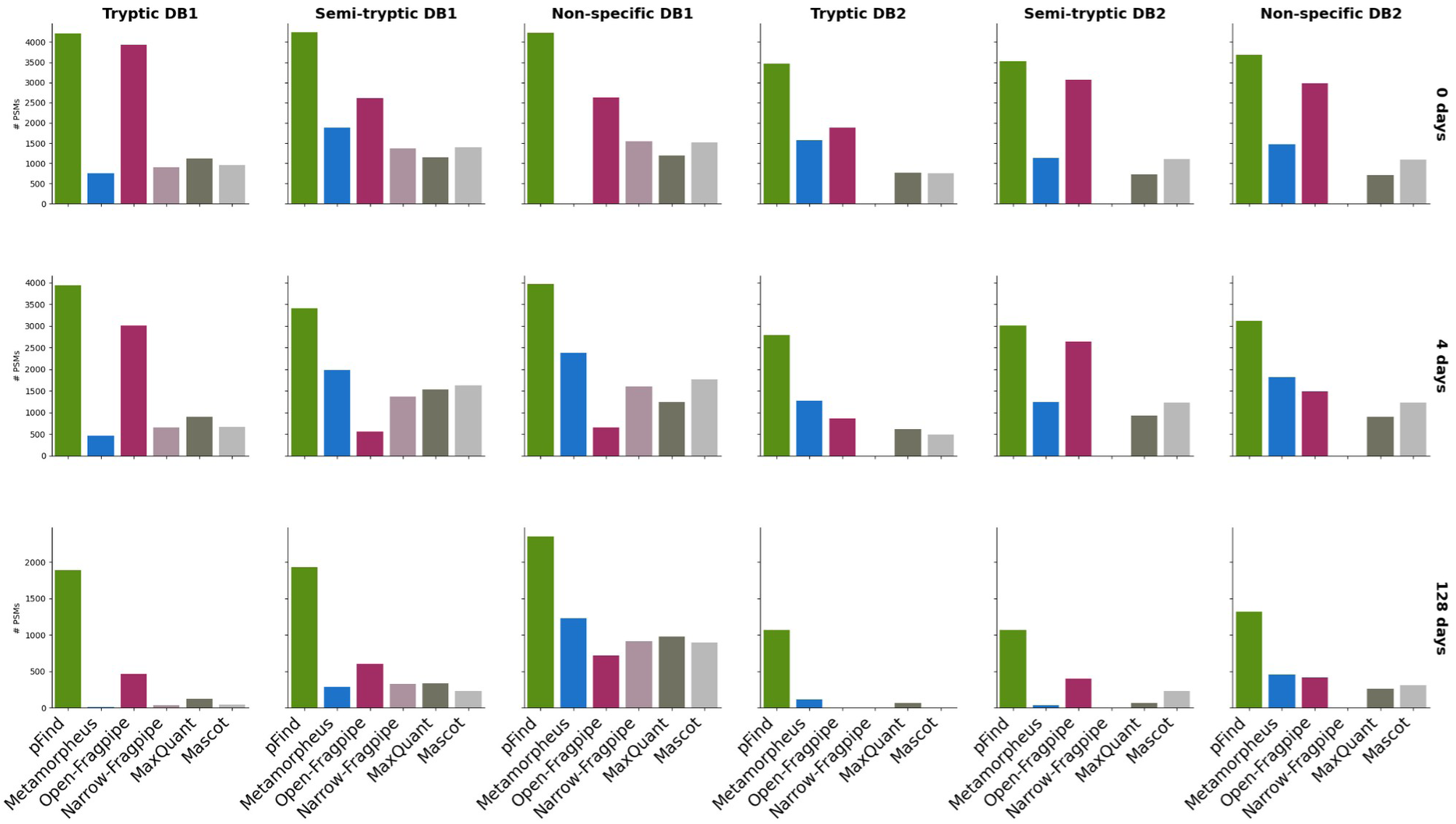
Number of PSMs at FDR 0.01 for each database search software, enzyme cleavage and database.

**Supplementary Figure S2.**
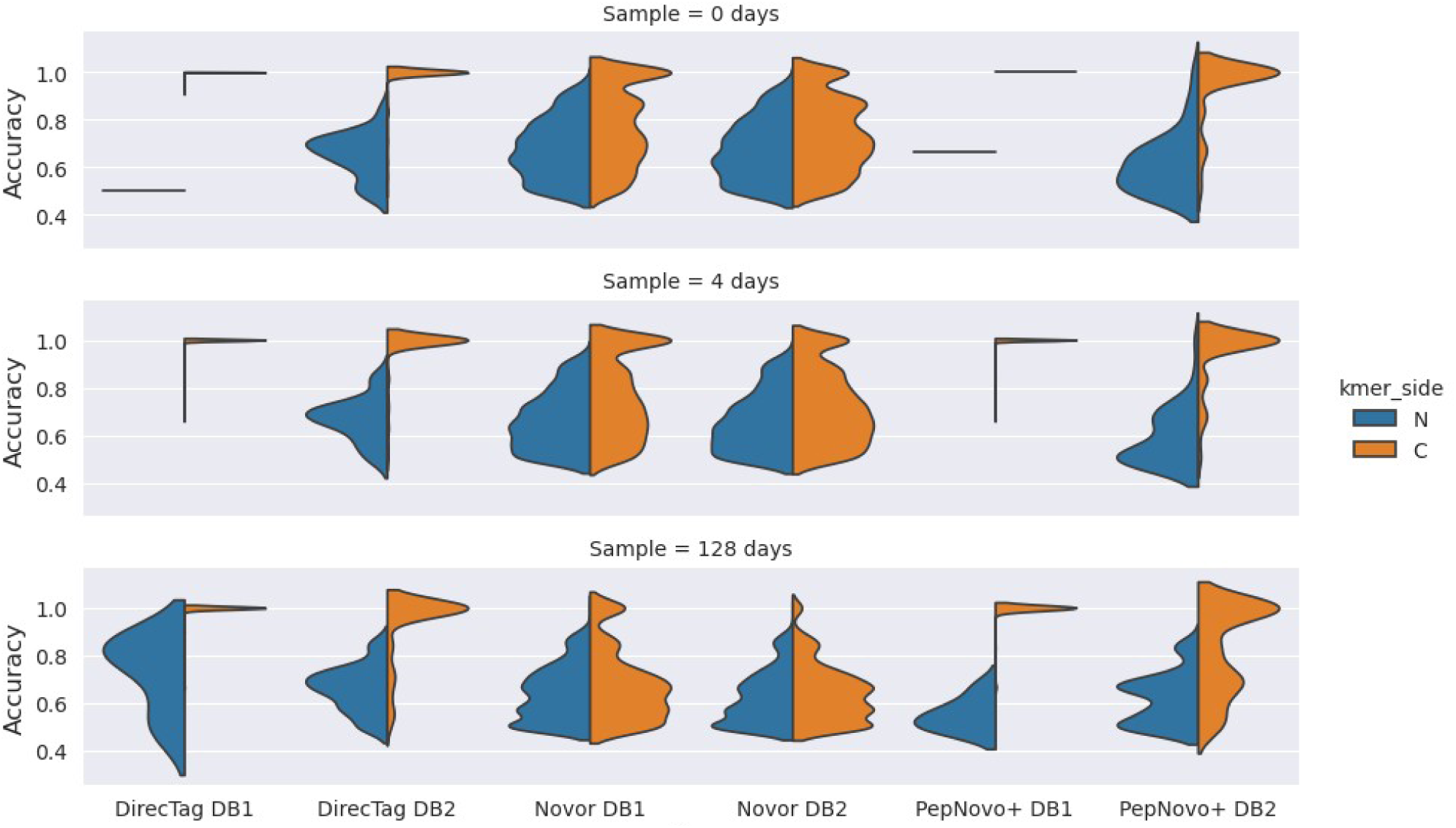
Accuracy distribution of *de novo* peptides aligned to the bovine β-lactoglobulin. In this plot, we chose the best alignment for each peptide, irrespective of whether it came from a N or C-terminal kmer seed alignment. We separate the accuracy of alignments coming from either a N-terminal (blue) or C-terminal (orange) seed k-mer.

**Supplementary Table S1.** Number of PSMs per run and software and at FDR 0.01 and 0.05.

| Digestion | Database | Engine | FDR 0.01 |  |  | FDR 0.05 |  |  |
| --- | --- | --- | --- | --- | --- | --- | --- | --- |
|  |  |  | 0 days | 4 days | 128 days | 0 days | 4 days | 128 days |
| Tryptic | DB1 | Mascot | 949 | 668 | 46 | 1076 | 753 | 49 |
|  |  | MaxQuant | 1119 | 902 | 123 | 1119 | 902 | 123 |
|  |  | Fragpipe-narrow | 891 | 648 | 31 | 917 | 725 | 40 |
|  |  | Metamorpheus | 748 | 466 | 6 | 1209 | 802 | 104 |
|  |  | Fragpipe-open | 3930 | 3004 | 464 | 4639 | 3949 | 605 |
|  |  | pFind-open | 4206 | 3938 | 1887 | 4666 | 5018 | 2258 |
|  | DB2 | Mascot | 757 | 484 | 0 | 869 | 560 | 0 |
|  |  | MaxQuant | 772 | 619 | 69 | 772 | 619 | 69 |
|  |  | Metamorpheus | 1578 | 1273 | 115 | 1967 | 1558 | 304 |
|  |  | Fragpipe-open | 1886 | 859 | 0 | 2790 | 1229 | 0 |
|  |  | pFind-open | 3468 | 2785 | 1069 | 4088 | 3738 | 1257 |
| Semi-tryptic | DB1 | Mascot | 1397 | 1625 | 231 | 1612 | 1836 | 280 |
|  |  | MaxQuant | 1153 | 1528 | 338 | 1292 | 1738 | 447 |
|  |  | Fragpipe-narrow | 1343 | 1096 | 299 | 1443 | 1252 | 381 |
|  |  | Metamorpheus | 1877 | 1983 | 289 | 2712 | 3064 | 1030 |
|  |  | Fragpipe-open | 2618 | 552 | 602 | 2874 | 644 | 1851 |
|  |  | pFind | 4246 | 3403 | 1927 | 4726 | 5212 | 2328 |
|  | DB2 | Mascot | 1109 | 1225 | 231 | 1344 | 1470 | 280 |
|  |  | MaxQuant | 719 | 931 | 66 | 853 | 1099 | 136 |
|  |  | Metamorpheus | 1127 | 1237 | 38 | 1845 | 2107 | 345 |
|  |  | Fragpipe-open | 3073 | 2636 | 401 | 3798 | 3358 | 472 |
|  |  | pFind | 3520 | 3006 | 1066 | 4188 | 4007 | 1305 |
| Non-specific | DB1 | Mascot | 1506 | 1757 | 893 | 1689 | 1991 | 1036 |
|  |  | MaxQuant | 1183 | 1241 | 974 | 1336 | 1422 | 1233 |
|  |  | Fragpipe-narrow | 1424 | 970 | 657 | 1530 | 1088 | 766 |
|  |  | Metamorpheus | 0 | 2382 | 1231 | 0 | 3067 | 2192 |
|  |  | Fragpipe-open | 2621 | 652 | 718 | 2863 | 687 | 1516 |
|  |  | pFind | 4221 | 3967 | 2351 | 4796 | 5482 | 3067 |
|  | DB2 | Mascot | 1093 | 1235 | 311 | 1377 | 1543 | 413 |
|  |  | MaxQuant | 710 | 900 | 264 | 852 | 1127 | 451 |
|  |  | Metamorpheus | 1467 | 1817 | 455 | 1791 | 2255 | 934 |
|  |  | Fragpipe-open | 2973 | 1493 | 417 | 3682 | 1839 | 666 |
|  |  | pFind | 3682 | 3123 | 1321 | 4283 | 4200 | 1662 |

Supplementary Table S2. Number of PSMs for each run and software and at FDR thresholds 0.01 and 0.05.

## Supplementary Files

**S1**. parameters_table.xlsx is a spreadsheet with all the search parameters for MaxQuant, Mascot, Metamorpheus, pFind and Fragpipe. These are derived directly from the parameter files used to set up each software and run. Variable parameters (enzyme or database, or related to open/narrow search in Fragpipe) are highlighted.

